# Directed evolution of a thermostable family-A DNA polymerase into a reverse transcriptase for 2’-modified RNA

**DOI:** 10.64898/2026.10.08.757645

**Authors:** Eloi Vincent, Clément Lopez, Rémi Sieskind, Isciane Commenge, Rafael Navaza, Ahmed Haouz, Marc Delarue

## Abstract

RNA-based therapeutics increasingly rely on nuclease-resistant xeno-nucleic acids (XNAs), but engineering polymerases capable of reading and amplifying these non-canonical substrates remains a major challenge. Natural DNA polymerases cannot process C2’ sugar modifications, and the complete structural determinants governing this limitation remain poorly understood, limiting rational design strategies.

In this study, we apply directed evolution methods to *Thermus thermophilus* DNA polymerase using a microfluidic implementation of the Compartmentalized Self-Replication (CSR). We identify an engineered variant, termed E5, capable of reverse transcribing C2’-modified RNA, a previously inaccessible activity for this enzyme. Next-generation sequencing allow us to track evolutionary trajectories across nine rounds of evolution. Through an integrated analysis combining statistical methods, protein language models, and crystallographic data, we reveal unknown epistatic networks controlling polymerase function. This large-scale experimental dataset (>196,000 thermostable enzyme variants) provides a robust basis for predicting sequence-to-function relationships, accelerating development of enzymes for aptamer discovery and synthetic biology applications.

## Introduction

Protein design now achieves near-atomic control of molecular folding^1^, but extending similar design principles to complex enzymatic mechanisms remains difficult, particularly for reactions on non-natural substrates. Nucleic-acid-processing enzymes are particularly attractive targets for such efforts, given their central role in emerging RNA-based therapeutics. Available enzymes are currently unable to efficiently process the chemically modified substrates required by these applications. Reverse transcriptases (RTs) are central to these applications, but most naturally occurring RTs are poorly thermostable and unable to efficiently read synthetic nucleic acid analogues called XNAs^2–4^. This limits their integration into numerous applications such as reading modified nucleic acid substrates in the selection of therapeutical aptamers during SELEX cycles^5,6^ or combining RT and PCR steps for the diagnosis of viral infections through quantitative reverse-transcription PCR^7^.

Directed evolution takes advantage of already thermostable DNA-dependent DNA polymerases^7^ as chassis for engineering RT activity, using technologies such as phage display ^8,9^, droplet-based screening such as CBL^10^ or self-selection processes like compartmentalized self-replication (CSR)^11^. The latter leverages a positive feedback loop where only polymerases capable of PCR amplifying their own gene under selective conditions are enriched over iterative rounds^11^. RT-CSR, a CSR variant in which polymerases must bypass a patch of ribonucleotides to amplify their own genes, has proven particularly powerful for generating thermostable RTs. Efforts have focused predominantly on archaeal family-B polymerases (such as from *Thermococcus kodakarensis*) and produced enzyme performing RT-PCR on more than 5 kb RNAs but without an exhaustive analysis of the mutational landscape leading to the selected unnatural activity^12,13^. In contrast, bacterial family-A polymerases can exhibit latent RT activity that is typically revealed only in the presence of Mn²⁺ ions^14^, usually with a decreased fidelity, and their potential as platforms for robust RT engineering remains unexplored by directed evolution. In particular, if the goal is to engineer a DNA polymerase capable of reverse transcribing RNA bearing 2’-O-methyl (2’OMe) modifications such as for the selection of nuclease-resistant aptamers, it remains unclear whether family-A DNA polymerases (PolA) offer a superior scaffold compared to family-B DNA polymerases (PolB). A schematic representation of the evolutionary relationship between PolA, PolB, RT and RNA-dependent RNA polymerases is presented in **Figure 1a**.

**Figure 1.**
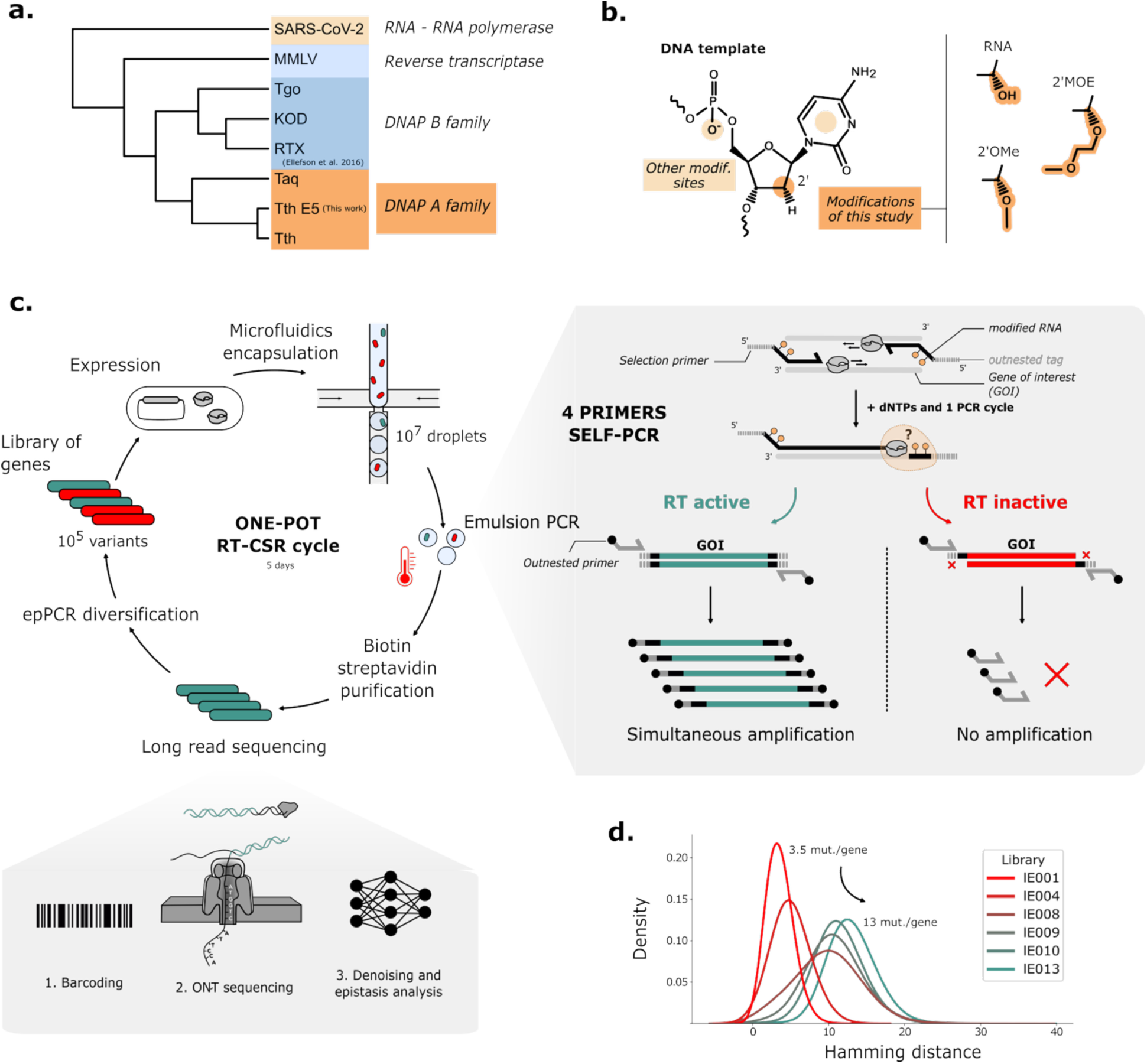
Directed evolution strategy for engineering thermostable reverse transcriptase (RT) activity in *Thermus thermophilus* DNA polymerase. **A.** (a) Phylogenetic relationships of major DNA and RNA polymerase families (DNAP, viral RdRp, and RT) highlighting the evolutionary distance between already engineered B-family DNAP and A-family DNAP engineered in this study. **(b)** Chemical structures of C2’-modified templates (RNA, 2’OMe, 2’MOE) used in this study. **(c)** RT-CSR cycle. A library of Tth DNA polymerase A variants is generated, expressed in *E. coli*, and compartmentalized using microfluidics for Reverse Transcription Compartmentalized Self-Replication (RT-CSR). Within droplets, a four-primer self-amplification reaction is performed, where gene of interest (GOI) flanking primers contain variable combinations of 2’MOE and RNA bases to probe RT activity. A second set of 5’-biotinylated primers, supplied in excess, hybridizes selectively to the outnested tag product upon successful reverse transcription. Active variants are isolated by streptavidin–biotin purification, recloned, and analysed using Oxford Nanopore long-read sequencing. **(d)** Distribution of amino acid substitutions relative to Tth PolA sequence in successive rounds of directed evolution.

Structural studies have identified residues responsible for polymerase ability to form a complex with a primer-template DNA duplex and with an incoming nucleotide in family-A DNA polymerases. They revealed the high similarity between *Thermus aquaticus* (Taq)^15^, *Bacillus stearothermophilus* (Bst)^16^, and bacteriophage T7^17^ polymerases structure and mechanism. Beyond structural characterization, large-scale sequence alignments have enabled the identification of conserved motifs across this family mediating interactions with DNA and the catalytic process (motifs A, B, and C)^18,19^. Integrating sequence and structural information has facilitated targeted engineering efforts to alter enzyme specificity. For instance, this approach enabled the conversion of Taq DNA polymerase into an RNA polymerase, by identifying key residues forming a “steric gate” within motif A (I616E, M or T and E617G)^20–22^ and motif B (F669A)^23^ that prevent incorporation of ribonucleotides and mutating them. A more elaborate steric gate was even identified in family-B DNA polymerases^24^. Comprehensive mutational analysis of motifs A and B^25,26^ further documented their functional plasticity. Building on this knowledge, efficient XNA reverse transcriptases were identified in family-B polymerases^10,27–29^, but natural family-A DNA polymerases remain to be engineered to acquire robust XNA reverse transcriptase activity.

Here we show that a microfluidic RT-CSR strategy coupled with next-generation sequencing is efficient to enhance the intrinsic RT-XNA activity of a *Thermus thermophilus* family-A DNA polymerase (Tth) for the challenging 2’-O-methoxyethyl (2’MOE) RNA reverse transcription. This modification, which differs from previously studied 2’OMe groups (**Figure 1b**), increases duplex melting temperature and shifts the helix from B-form to the A-form^30^. It thereby imposes stringent structural constraints on polymerase function. The evolved enzyme called E5 operate without Mn²⁺ stimulation, display activity on highly modified templates, and the final and intermediate libraries of selected mutants can be investigated using next-generation sequencing and crystallographic structures to understand exact evolutionary trajectories. This also provides high-quality datasets for training AI-methods currently developed for enzyme design. In the long term, this study provides a general route towards more robust and versatile reverse transcriptases for RNA and XNA biotechnology.

## Results

### Microfluidics-based RT-CSR

The microfluidics-based RT-CSR was implemented so as to enable efficient engineering of a family-A DNA polymerase. It overcomes limitations of previous RT-CSR such as the complex two PCR steps and the high emulsion polydispersity. Mutants libraries were transformed into *E. Coli* bacteria to express the mutant polymerases. They were then encapsulated with dNTPs, selective primers and 5’-biotinylated primers into water-in-oil droplets using flow focusing microfluidic chip and following Poissonian distribution with a mean-value λ=0.1. The obtained emulsion was submitted to PCR cycles and, after emulsion break, the amplicons mixed in the resulting aqueous phase were purified and recloned (**Figure 1c**). While microfluidics led to reduced final product concentration, the high monodispersity provides superior control over reaction conditions and minimized co-encapsulation artefacts that can disrupt enzyme evolution^31^. Method performance was validated in a control selection, where a mixture of active and inactive variants yielded selective enrichment of only the active clones **(Supplementary Figure 1)**. The stability and monodispersity of such emulsions ensured droplet-to-droplet similar selection conditions and increased variant enrichment reproducibility. To further improve the workflow, we incorporated a biotin-streptavidin purification of the emulsion PCR products and decreased bacterial growth time. These adjustments markedly reduced contamination from undesired sequences (e.g., parental DNA or sequences with premature stop codons yielding truncated proteins) **(Supplementary Figure 2)** and improved the enrichment of functional RT polymerase variants. Sanger sequencing analyses of hundreds of clones revealed a progressive accumulation of non-silent mutations during evolution, increasing from an average of 3.2 ± 1.5 per gene at early cycles to 13 ± 2.9 per gene at completion (**Figure 1d**).

### Evolution of Tth polymerase activity across evolution rounds

Once the protocol was established, we initiated RT-CSR experiments using the wild-type (WT) Tth PolA sequence, in contrast to previous engineering efforts that relied on pre-engineered enzymes as starting points^32^. We conducted nine rounds of selection with the intercalation of four error-prone PCR diversification cycles and gradual increases in selection stringency (**Figure 2a**). From the third cycle onward, the mutagenized region was expanded from 500 to 1,000 base pairs to encompass all the catalytic motifs (TGR, A, B, 6 and C defined in the literature^18,19,33^) and not only the region around motifs A and B initially targeted. Beneficial mutations were accumulated and not substituted. In the first three rounds, mutations such as A607T, I616F, F751Y, and M753T appeared with a frequency exceeding 20%. Later rounds (6-9) were characterized by specific enrichment of V739D and M749I, and the addition of L505P and L554Q (**Figure 2a**). The I616F mutation, located next to the steric gate position of PolA (E617), persisted throughout the experiment and is known as a key determinant of substrate specificity in this polymerase family^34^.

**Figure 2.**
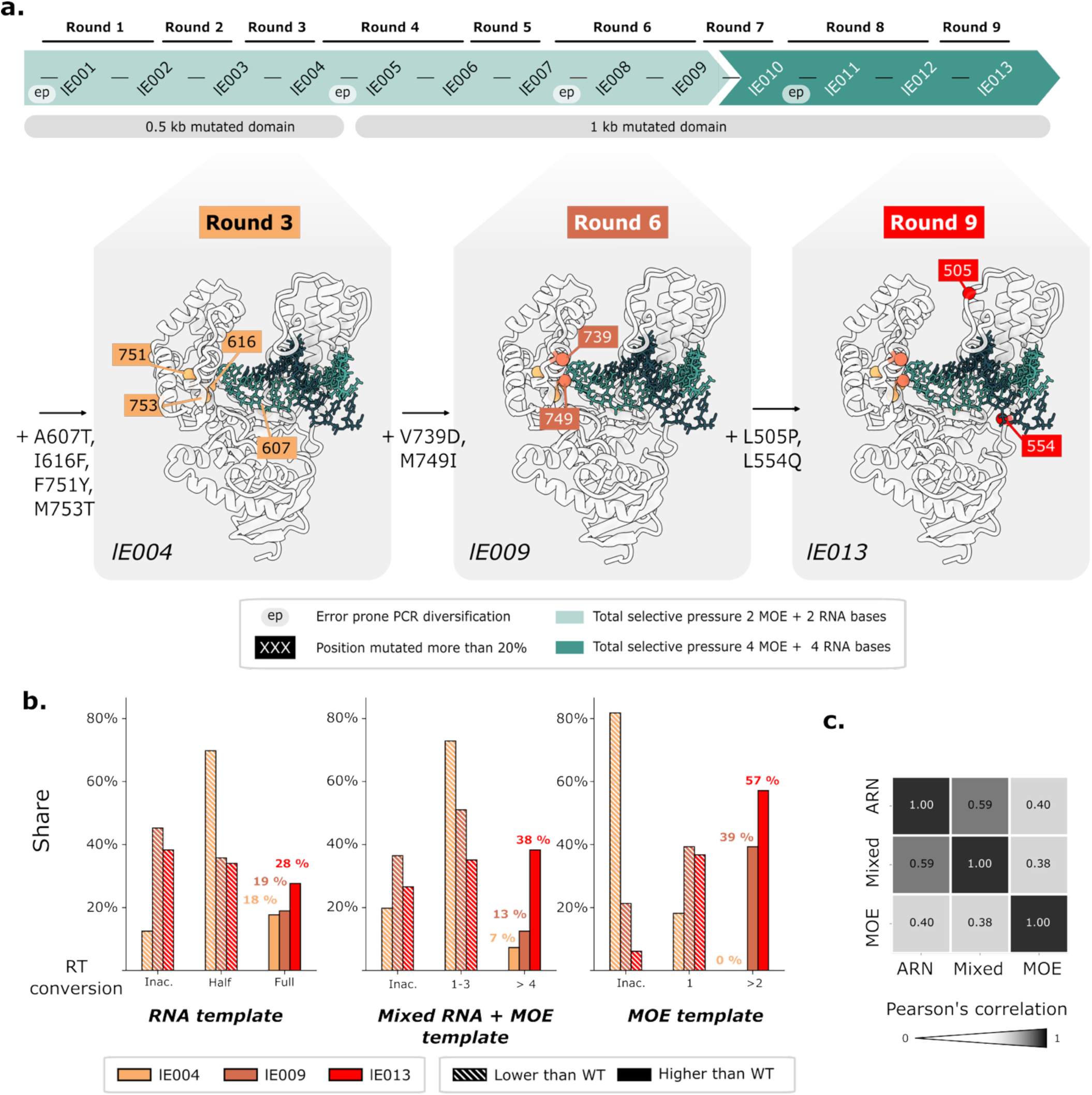
Experimental evolution and functional characterization of thermostable reverse transcriptase (RT) libraries. **(a)** Nine rounds of directed evolution interspersed by four error-prone diversifications were performed using selective primers containing both 2’MOE and RNA nucleotides. The initial selection targeted a 500 bp domain, which was extended to 1000 bp after the third round to encompass the three catalytic motifs (A, B, and C). The most mutated positions were mapped onto the crystal structure of WT Tth PolA. Mutations occurring in more than 20% of the final population (L505P, L554Q, A607T, I616F, V739D, M749I, F751Y, and M753T) are highlighted. **(b)** RT activity was monitored across evolutionary rounds by screening 96 individual randomly chosen clones against various templates (RNA, 2’MOE, and mixed RNA/2’MOE). In the final library, 57% of active clones outperformed the WT in the most stringent 2’MOE conversion assay. **(c)** Pearson’s correlation coefficients between activities on different substrates across all libraries indicate that performance on mixed RNA+MOE and RNA templates is strongly correlated. This suggests that improved activity on one substrate translates to enhanced activity on other C2’-modified templates.

We showed that overall the library activity against chemically modified substrates improved progressively through the course of evolution. Three forms of modified substrates were evaluated: consecutive RNA bases, consecutive 2’MOE bases, and a mixed RNA: 2’MOE (1:1) substrate analogous to the one used for selection. Screening of 96 random clones from three libraries at rounds 3, 6, and 9 indicated systematic gains in XNA-RT performance. More than 80% of variants were inactive with the highly stringent full 2’MOE substrate at round 3, while inactivity fell to just 5% at round 9. Notably, 57% of random clones at round 9 outperformed WT polymerase. 38% of variants surpassed the WT for 2’MOE/RNA mixed substrate, and 28% did so for the all-RNA substrate at the end of the evolution (**Figure 2b**). Activities measured on mixed and RNA template were highly correlated (Pearson’s coefficient = 0.59), and variants displaying activity for the most complex substrates (full 2’MOE) also performed well on simpler ones (RNA and mixed) **(Figure 2c)**.

### Variant E5 outperforms WT polymerase

By the end of round 9, a polymerase variant, named E5, was isolated and tested for activity. It displayed markedly enhanced activity on multiple substrates compared to the WT enzyme. Primer extension assays were performed with purified enzymes **(Supplementary Figure 3)** using DNA substrates with one modified base (RNA, 2ʹOMe or 2ʹMOE). Conversion kinetics were then analyzed by capillary electrophoresis. Across all tested substrates, E5 exhibited higher conversion efficiencies than the parental polymerase, with the most pronounced improvement observed for the challenging 2’MOE-modified substrate. Whereas the WT enzyme failed to bypass the 2’MOE base even after 1 h of incubation (stalling one base before the modification or falling from the template), E5 converted approximately 60% of the 2’MOE substrate within 1 h and 20% within just 2 min. This result indicates that directed evolution improved substantially the parental polymerase with the desired noncanonical activity. The blocking of the WT supports the notion that a very substantial evolutionary barrier was crossed to efficiently process 2’MOE modifications **(Figure 3a)**. Fitting the time courses to a single-exponential model yielded apparent rate constants *k_obs_*, which quantify this gain in catalytic efficiency. E5 displayed approximately fourfold higher *k_obs_* than the WT enzyme on both DNA and RNA substrates, and a 81-fold increase in *k_obs_* on the 2ʹOMe substrate. Notably, the E5 evolved enzyme exhibited comparable *k_obs_* values for 2ʹOMe and RNA single base bypass (respectively 1.7s^-1^ versus 1.2s^-1^), whereas the WT enzyme showed substantially reduced activity on the 2ʹOMe-modified substrate (respectively 0.28s^-1^ versus 0.021s^-1^). **(Figure 3b)**.

**Figure 3.**
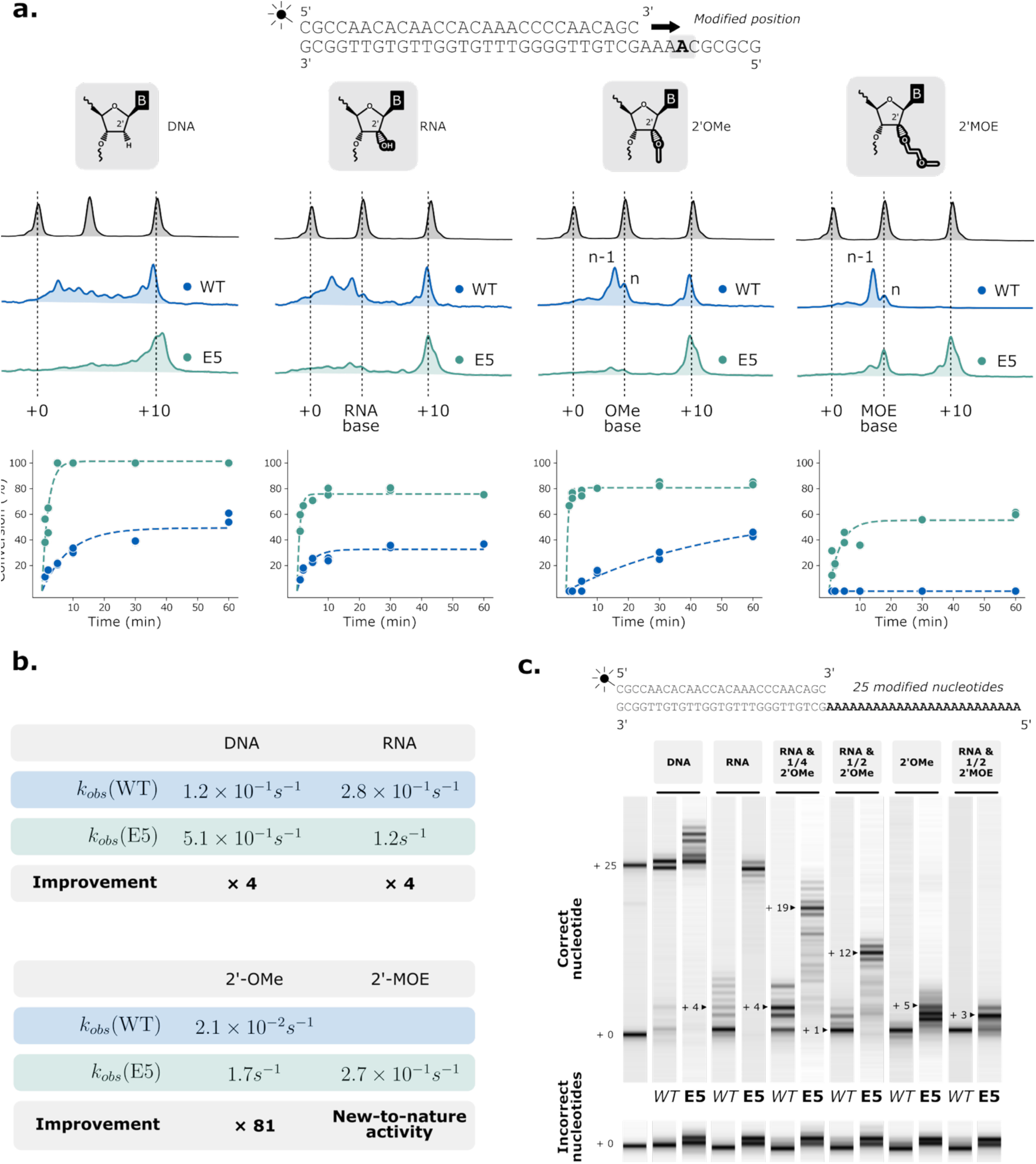
Functional characterisation of the evolved polymerase variant E5. **(a)** Capillary electrophoresis of single-base RT assays. Primer extension kinetics are measured for the WT (blue) and evolved (E5, green) polymerases. The DNA templates contain one RNA, 2ʹOMe or 2ʹMOE residue at position +4. **Top**: Capillary electrophoresis traces of 60 min reaction products show unextended primers (+0), fully extended products (+10), or intermediates stalled at the modified base. **Bottom**: Conversion to full-length product measured as the fraction of fully extended versus stalled products from integrated peak areas over time (duplicate reactions per time point). Dashed lines represent best fits to a single-exponential model used to extract apparent rate constants (*k_obs_*). **(b)** Kinetic parameters, showing *k_obs_* values and relative catalytic efficiency enhancements for WT and E5 polymerases. E5 displayed approximately fourfold higher *k_obs_* on DNA and RNA substrates, an 81-fold improvement on the 2ʹOMe substrate, and a newly acquired activity toward the non-natural 2ʹMOE substrate. The WT enzyme fails to extend beyond the 2’MOE modified position even after 1 h, whereas E5 reaches 60 % conversion after 1 h and 20 % after 120 s, with the WT remaining stalled one nucleotide before the modification. **(c)** A fidelity assay was performed using poly(A) templates composed of consecutive modified bases, comparing the incorporation of the correct nucleotide (dTTP) with that of incorrect nucleotides (dATP, dGTP, or dCTP). The evolved variant E5 incorporated the correct nucleotide more efficiently than the WT across all fully-modified tested substrates: RNA (+4 vs +25), RNA with 25% 2ʹOMe (+4 vs +19), RNA with 50% 2’OMe (+1 vs +12), 2’OMe (+1 vs +5) and RNA with 50% 2ʹMOE templates (+1 vs +3). For incorrect nucleotides, E5 incorporated one or two nucleotides and then stops.

E5 activity was further assessed on diverse templates including homopolymeric substrates (DNA, RNA, or 2’OMe) and RNA sequences containing 25% 2’OMe, 50% 2’OMe, or 50% 2’MOE modifications. On unmodified RNA templates, E5 achieved complete RT conversion in 30 mn whereas the WT stalled after 4 bases. For mixed RNA/2’OMe templates, E5 successfully extended across 19 consecutive nucleotides (25% modification) or 12 nucleotides (50% modification), whereas WT stalled after 4 or 1 nucleotides, respectively. On fully 2’OMe templates, E5 bypassed 5 consecutive modified nucleotides compared to only 1 for WT. For mixed RNA/2’MOE templates, E5 traversed 3 modified positions versus 1 for WT, demonstrating substantially enhanced processivity across consecutive C2’ modifications **(Figure 3c)**. Fidelity assessment using incorrect nucleotides in front of the poly(A) template revealed a marked reduction in fidelity compared to WT **(Figure 3c)**.

Mutant E5 harbors 13 non-synonymous mutations (L505P, P552S, L554Q, A607T, I616F, Y698N, E710V, V739D, V742A, M749I, F751Y, M753T, A796E). Eight of the E5 mutations correspond to the most highly enriched positions in the final library, whereas the remaining five substitutions rank between the 251^st^ and 1115^th^ most frequent amino-acid changes, indicating a combination of strongly selected mutations and lower-frequency substitutions that contribute to the overall E5 fitness (**Figure 4**). Seven mutations appear to increase the polarity of the catalytic pocket. Overall, mutations cluster near the DNA-binding site and the catalytic core but positions 552, 710 and 796 are the most solvent-exposed. Among the mutated residues, the distances to the active site range from 9.8 to 32.7 Å, with I616 and M753 forming the closest mutations to the C3ʹ primer atom (9.8 and 15.8 Å, respectively). Several other positions remain over 25 Å away from the ligand. Notably, no mutation at position E617 (the steric gate) was found, while it often undergoes bulky-to-small residue substitution according to literature^23^ but its immediate neighbor I616 was found mutated as in other related studies ^20,22^. BLOSUM scoring revealed that most mutations in E5 (8 out of 13) are non-conservative, indicated by negative scores. It suggests that the acquisition of novel substrate specificity require substantial destabilizing changes, as already shown in other previous evolution campaigns^35^ (**Figure 4a**). Most of these mutations cluster in the fingers and thumb domains (10/13), as well as within conserved motifs 6 and A (5/13) (**Figure 4b**).

**Figure 4.**
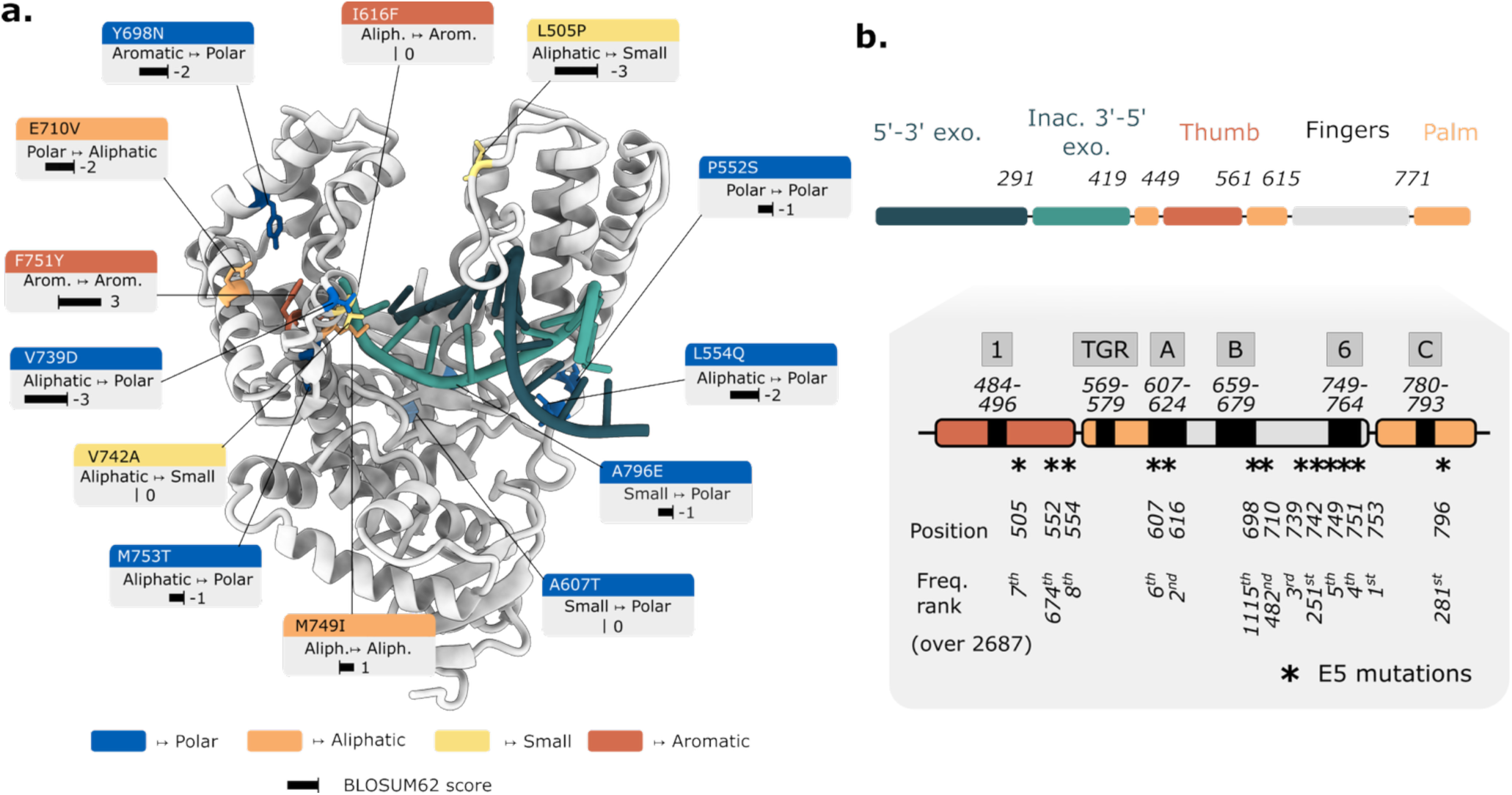
Structural analysis of E5 mutations driving enhanced reverse-transcriptase activity. **(a)** The E5 variant carries 13 amino-acid substitutions (L505P, P552S, L554Q, A607T, I616F, Y698N, E710V, V739D, V742A, M749I, F751Y, M753T, A796E), most of which are non-conservative according to BLOSUM scoring (score < 0), consistent with the radical replacements often observed in DNA polymerases engineered for XNA or RT activity^73^. Eight of these substitutions increase side-chain polarity near the catalytic pocket. The mutations are indicated by a color code: Polar (blue), Aliphatic (orange), Small (yellow), Aromatic (brown), where these classes of amino-acids are defined according to Dayhoff et al. ^74^. **(b)** The 13 substitutions in E5 are distributed across the palm (3/13), fingers (7/13), and thumb (3/13) domains of the polymerase. Motifs A and 6 (corresponding to the N-terminal portion of helix Q in the fingers domain) are particularly affected. Eight of the E5 mutations correspond to the most highly enriched positions in the final selected library, whereas the remaining five substitutions rank between the 251^st^ and 1115^th^ most frequent amino-acid changes, indicating a combination of strongly selected mutations and lower-frequency substitutions that contribute to the overall E5 fitness.

### Temperature dependence of RT activity

The RT-CSR directed evolution campaign developed in this study relies on the enzyme’s ability to perform droplet-based PCR cycles. This PCR requires high thermal resistance, so we tested the acquisition of any improved thermostability by the E5 variant. Two reverse transcriptase assays were conducted: one involving progressive denaturation of the enzyme at 90 °C followed by RT activity measurement, and another testing a range of elongation temperatures. These experiments demonstrate that the evolved variant is RT active over a broad temperature range and exhibits higher reverse transcription activity than WT. Complete complementary DNA synthesis of a 25-nucleotide RNA substrate was achieved from 60 to 90 °C, whereas the parental enzyme is nearly inactive above 80 °C and never reaches full conversion **(Supplementary Figure 5)**. In denaturation assays, the thermostability of E5 is substantial. The WT enzyme has low basal activity for all incubation times, while the evolved E5 variant remains RT active even after 1.5 hours at 90 °C **(Supplementary Figure 6).** Differential scanning calorimetry analysis revealed similar thermostability for WT and E5, with melting temperatures (Tm) of 95.02 ± 0.014 °C and 92.80 ± 0.024 °C, respectively. The marginal 2.2 °C reduction in E5 Tm suggests that enhanced reverse transcriptase activity was achieved without compromising global protein stability.

### Structural investigation for poor WT 2’MOE RT activity

To understand how these mutations facilitate improved RT activity, we crystallized the WT Tth family-A DNA polymerase in presence of a 2’MOE modified template. The primer-template duplex had the same sequence as the one used by Waksman *et al.*^15^, who successfully crystallized Taq DNA PolA with dsDNA. The first templating nucleotide (n) was substituted by an RNA followed by a 2’MOE modified nucleotide (n+1) to mimic our selection substrate. Strikingly, the X-ray structure shows a DNA polymerase binding the XNA-containing duplex in the opposite direction, where the blunt-ended 3ʹ-OH of the template strand, rather than that of the primer, occupies the position normally used for extension in the enzyme. We can thus deduce that the single-stranded modified template is excluded from the active site and that this prevents any RT activity (**Figure 5b**).

**Figure 5.**
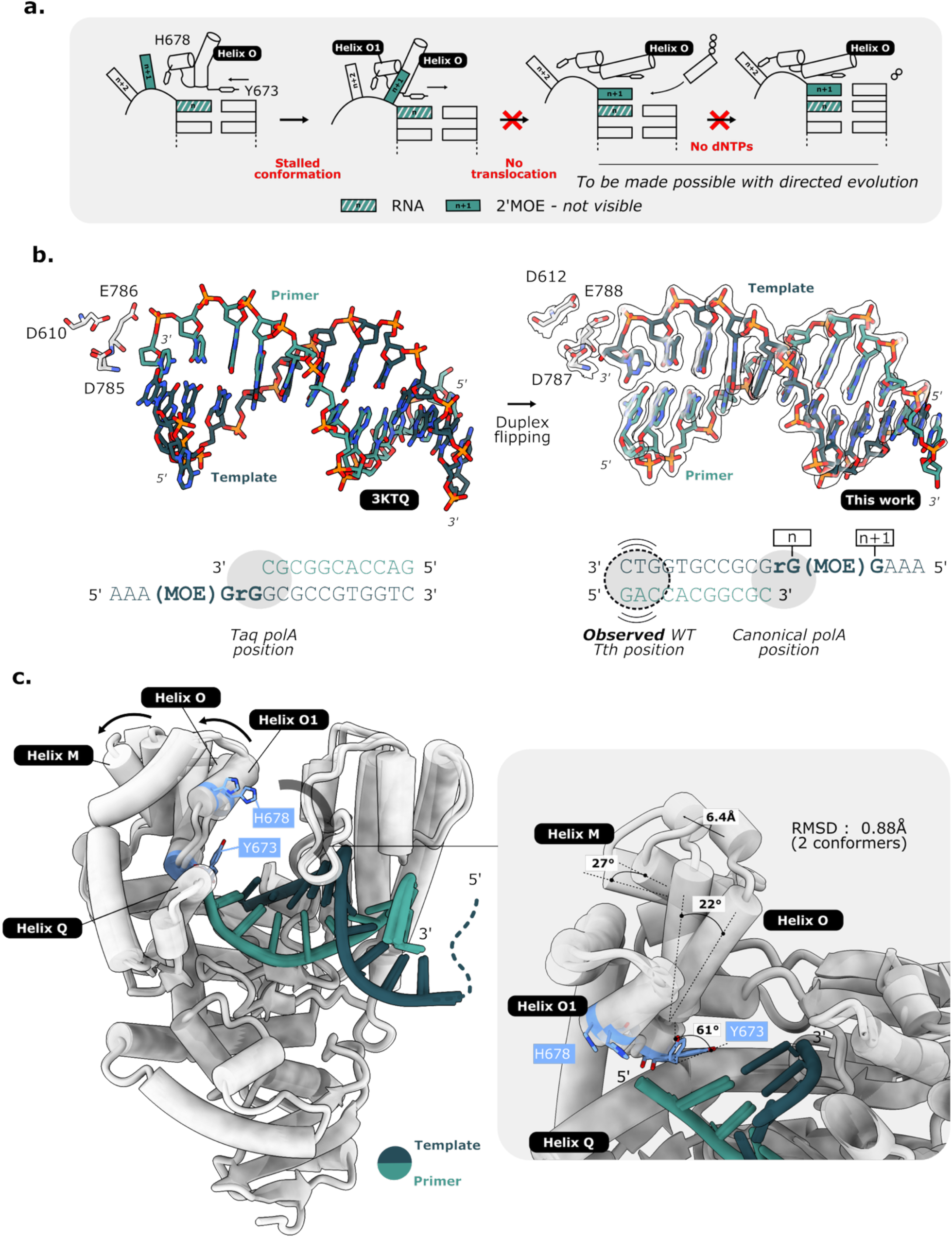
Structural and mechanistic basis for blocked RT activity by *T. thermophilus* PolA WT on a modified template. **(a)** Nucleotide incorporation by family-A polymerases is followed by translocation of the primer-template duplex. Histidine 678 stabilizes the next templating base (n + 1) via base stacking, while helix O orchestrates a push-pull motion of tyrosine 673 that checks the new base pair geometry. Following translocation, the correctly paired incoming nucleotide is positioned for phosphodiester bond formation. Blocked translocation prevents polymerization from proceeding as shown in ^36,37^. **(b)** The 1.9 Å crystal structure of the WT Tth enzyme bound to a duplex containing a modified 2ʹMOE residue shows that the polymerase fails to accommodate the modified template strand in a productive orientation. Instead, the duplex is captured in an inverted configuration in which the 3ʹ-OH of the template, rather than that of the primer, occupies the position normally used for extension, leading to a stalled RT activity. The obtained structure (right) is compared with the expected configuration for this polymerase A family (PDB:3KTQ, left). **(c)** The observed conformations prevents correct positioning of the next nucleotide as there is no templating base in position. The two main residues involved in translocation (Y673 and H678) are highlighted in pale blue. The resulting perturbation is associated with substantial flexibility of the fingers domain (approximately 6 Å displacement) and a 61° rotation of side-chain of the conserved tyrosine 673 residue, which cannot efficiently stack against the templating base.

The presence of two conformationally distinct molecules in the asymmetric unit provides two snapshots of Tth polymerase structure, highlighting pronounced mobility of helices M, N, and O within the fingers subdomain. Previous work demonstrated that nucleotide incorporation in PolA requires first a template translocation via a push-pull mechanism involving the highly conserved tyrosine 673, as well as a coordinated movement of helices O and O1 in the fingers subdomain, especially of residue H678 responsible for positioning the n+1 base, before binding of the incoming nucleotide^36,37^ (**Figure 5a**). In the Tth polymerase structure, both helices M and O are tilted by more than 20° in the two distinct forms. Strikingly, in one conformation, the conserved tyrosine 673 is flipped by 61°, and cannot fulfill its push-pull role. The second important residue involved in translocation, H678, is less destabilized as helix O1 stays more stable but can’t stack with the n+1 templating base. The two conformations differ by over 6 Å at the tip of the fingers domain (near helix N) but display an overall RMSD of 0.88 Å. (**Figure 5c**). This potential mobility of helix O mobility was seen by molecular dynamics for Bst polymerase in^38^ and measured by FRET^39^ but has not been captured by X-ray crystallography yet.

In the absence of an x-ray structure, AlphaFold3^40^ predictions were generated for WT and evolved E5 polymerases in complex with an RNA duplex bearing a 2’MOE modification at the first templating position and an incoming nucleotide. The most frequently observed conformation (present in 4 of 5 models) is shown for each enzyme in **Supplementary Figure 4**. In both cases, the RNA duplex and incoming nucleotide were positioned coherently with proper base pairing and high confidence scores (pTM > 0.9, ipTM > 0.7). Motif B (comprising helices O and O1) adopted markedly different conformations: in WT, motif B remained in an open conformation, whereas in E5 it adopted a closed state. This closed conformation in E5 enabled proper positioning of the n+1 2’MOE-modified templating nucleotide and incoming nucleotide. In WT, the incoming nucleotide mis-paired with the n+2 base, positioning the alpha phosphate 8.4 Å from the 3’OH site incompatible with catalysis. In contrast, E5 maintained a phosphate/3’-OH distance of around 4.9 Å. These structural differences suggest that directed evolution enabled proper translocation of the modified template and facilitates helix O closure as described in **Figure 5a**. The analysis of the most mutated region in E5, helix Q (motif 6), revealed additional mechanistic insights. The F751Y mutation, which emerged in early evolution rounds, improved accommodation of the 2’MOE aliphatic chain. Helix Q formed substantially more interactions with the modified template in E5 compared to WT.

The X-ray crystallographic structure and AlphaFold3 predictions illustrate how additional mutations are needed for promoting a correct positioning of the modified duplex in the polymerase active site and for reverse-transcription. For both structures, the fingers domain exhibits notable mobility in response to the resulting inverted conformation^41^, highlighting its key role in stabilizing and sensing the template strand.

### Mutational landscape and structural implications

High-throughput sequencing of six evolving libraries can be used to trace the origins of the experimentally observed reverse transcriptase activity as shown previously. The sequences are easily aligned and the multiple sequence alignment (MSA) reveals the following features. Several positions were strongly and recurrently mutated within the targeted and expected catalytic motifs (A, B, 6 and C defined in the literature^18,19,33^), such as positions 616, 751, and 753. However, mutational hotspots were distributed in an unexpectedly complex pattern. Prior to expanding the mutagenized zone from 500 to 1,000 base pairs, mutations primarily occurred at positions 607 and 616 (beta strand or alpha-helix of motif A), and at positions 749, 751, and 753 (beginning of helix Q of motif 6). Following the expansion of the mutated region and maintenance of selective pressure, after round 3 until round 6, additional positions became enriched, with a lot returning to baseline upon further rounds of selection and increase in selective pressure, like positions 659, 666, 696, 718, 732, 738, and 771. Some positions, such as 739, continued to show increasing mutation rate even as selective pressure intensified. Others, including positions in the palm region (776, 796, 807), fingers (638, 640, 648, 655, 667, 696, 692, 716), and thumb domains (505, 509, 518, 526, 531, 554), emerged only under higher selective pressure. Positional enrichment and depletion cycles were also seen, such as for positions 520, 669 and 678. Highly mutated positions were rarely observed in beta-turn-beta motifs such as TGR, C, or the region spanning positions 720-730 (**Figure 6a**). As expected, the E5 variant harbors all the most strongly enriched mutations, those exceeding 20% frequency in the library and standing out prominently**. (Supplementary Figure 7)**

**Figure 6.**
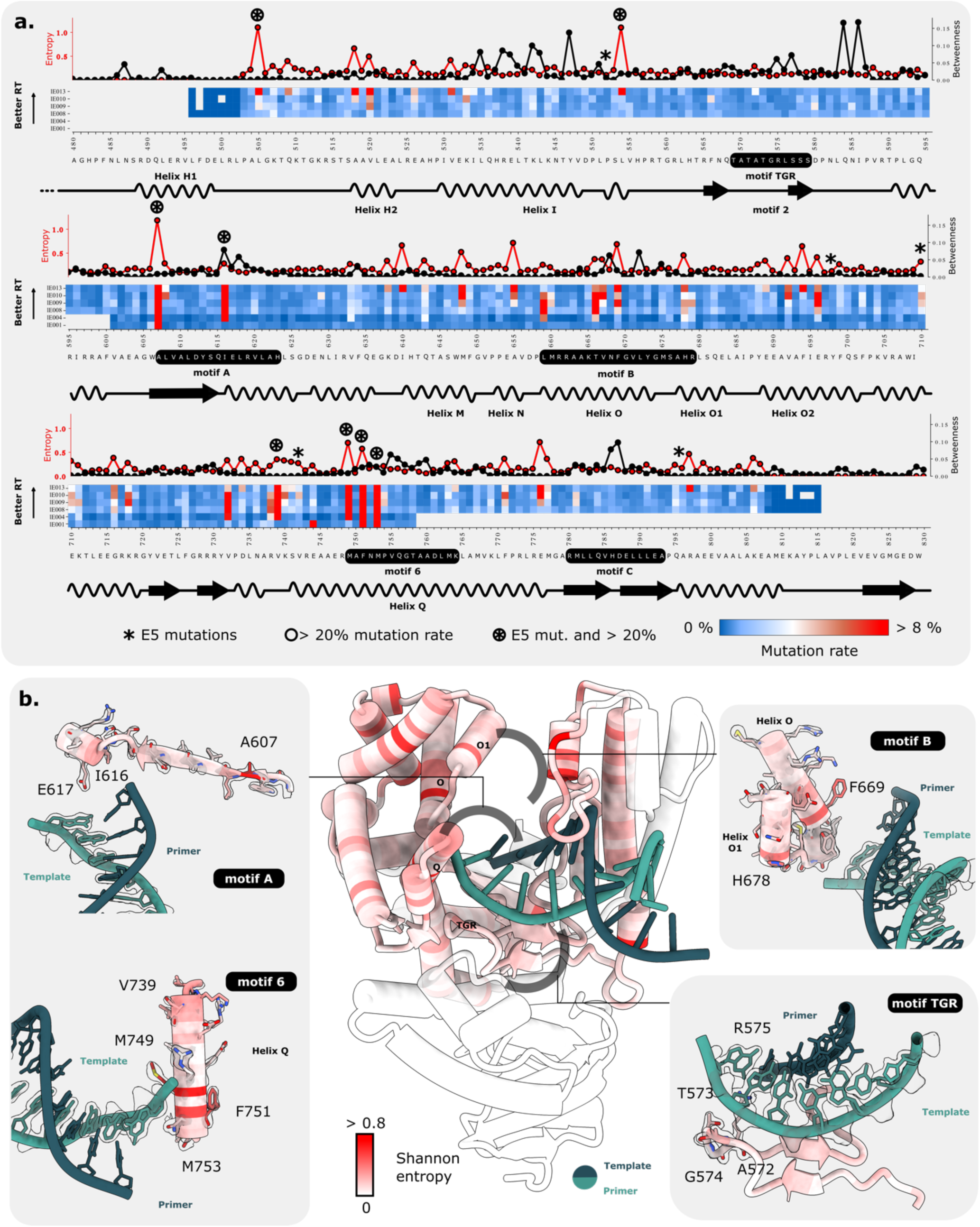
Single residue mutational hotspots underlying adaptation of Tth PolA during directed evolution. **(a)** Heatmap of mutational rates across sequence positions and evolutionary rounds, with Shannon entropy (red, left scale) and network betweenness centrality (black, right scale). Nanopore sequencing reveals mutational hotspots that do not correspond to residues with the highest network betweenness centrality, indicating no major changes in the structural amino acids network during evolution. Several positions show transient enrichment at low selection pressure but are subsequently depleted (for example, residues 718 and 771), whereas others become enriched only under increased selective pressure (residues 505, 531, and 554), mainly located within the thumb domain. The motif C mandatory for catalysis is barely mutated. **(b)** Per-residue Shannon entropy is mapped onto the polymerase structure and highlights how amino acid diversity per position changes across evolutionary rounds. Motifs A, 6, and B act as structural platforms facilitating acquisition of reverse transcriptase activity, while mutations within TGR and C motifs, despite their proximity to the template-binding region, tend to destabilize the enzyme.

Amino acid diversity (i.e the mutational tolerance) at each site was quantified using Shannon entropy and compared with centrality betweenness widely used in the study of protein interaction networks to quantify the connectivity role of some positions^42,43^ (**Figure 6a**). We observed that positions with the highest entropy did not correspond to inter-subunit communication hubs (higher betweenness). For instance, in the beginning of the motif B, the least-mutated position correlated with the highest betweenness centrality. Contrary to motif B, in motif A, the I616F mutation proved to be functionally and structurally critical as both Shannon’s entropy and betweenness centrality were consistently high at this position. Entropy did not solely reflect mutation frequency, as certain positions exhibit high mutation rates (I616 or F753) yet low entropy.

Projecting entropy hotspots onto the WT crystal structure facilitated interpretation (**Figure 6b**), revealing structural implications near catalytic motifs and highlighting that proximity to the template alone does not determine mutation likelihood. For instance, the TGR motif lies adjacent to the template but did not acquire significant mutations. Nevertheless, for motif B, aromatic amino acids F669 and H678, which are known to be important to interact with DNA and enable template translocation, were mutated. The proximity of position I616 in motif A to the nucleotide binding site suggests that the acquisition of reverse transcription activity requires repositioning of the incoming nucleotide in addition to the acquisition of the ability to translocate the modified template.

### Epistatic coupling analysis reveals functional networks

Single-point mutation analysis is informative for pinpointing positions responsive to selective pressure. However, this approach does not capture epistasis, the functional interactions between mutations. By implementing Nanopore sequencing and a robust variant-denoising pipeline, we obtained full-length sequences with every mutation for each variant. MSA for each evolutionary step enabled inference of functional coupling networks using a semi-physical Potts model based on column correlations in the MSA (mfDCA)^44^. This model provides an energy-based description inferred from the MSA, from which a probability of observing a natural or synthetic sequence can be calculated. The probability of a given sequence (*a*_1_, *a*_2_, …, *a_L_*) is expressed as an exponential function of a Hamiltonian (or statistical energy) parameterized by local fields ℎ*_i_*(*a_i_*), which reflects the frequency of each amino acid at each position, and pairwise couplings *J_i_*_j_\**a_i_*, *a*_j_+ that capture interactions between amino acids at two positions. Highly functionally coupled positions tend to change simultaneously. Here we report the first detailed application of this strategy to polymerase evolution for unnatural activity acquisition (**Figure 7a**). Direct information scores (DI) aggregates couplings from the Potts model and allows identification of high betweenness nodes.

**Figure 7.**
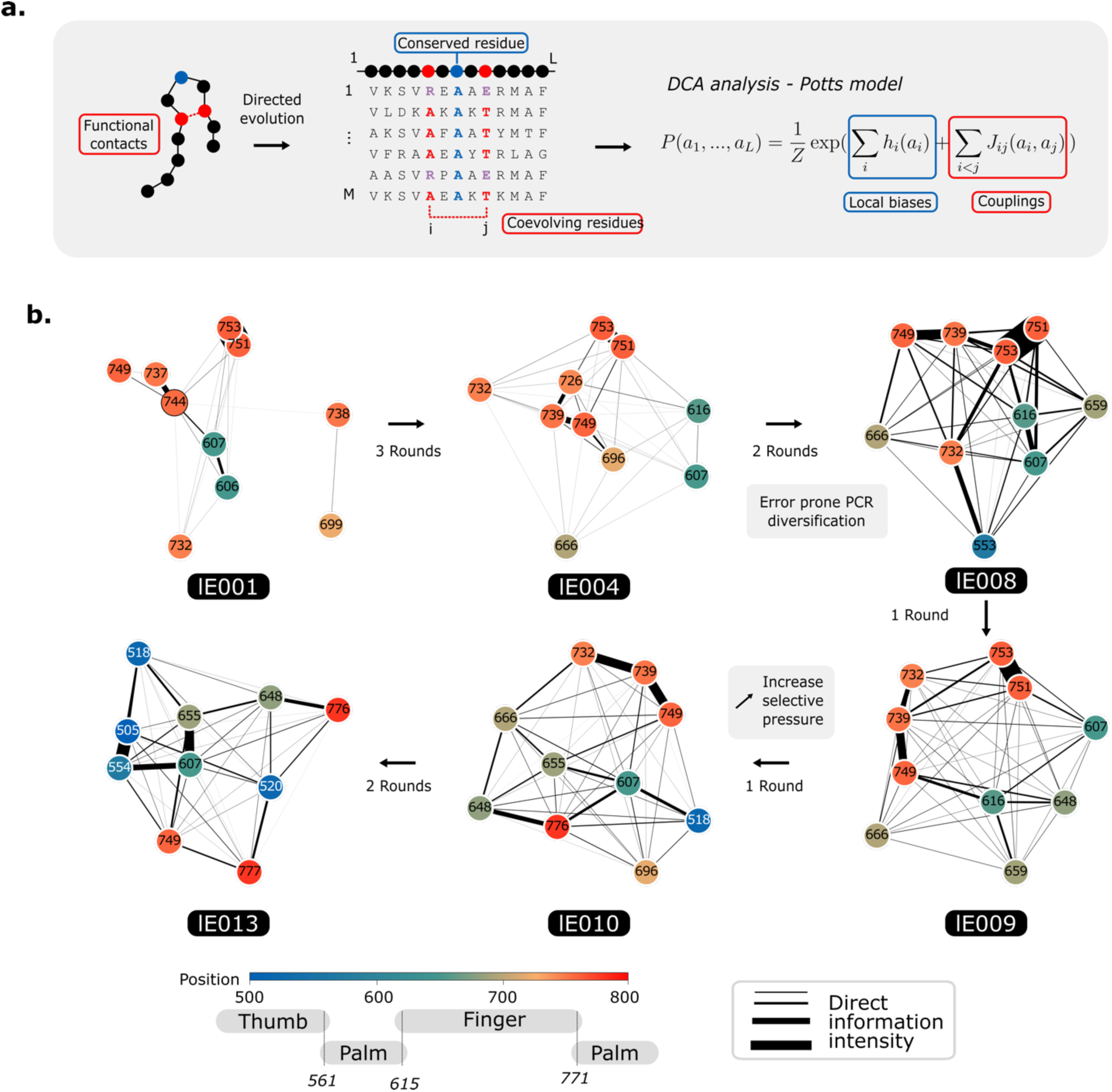
Coevolutionary constraints and residue coupling dynamics during polymerase evolution. **(a)** Evolutionary constraints such as those structuring the active site can be revealed by an MSA of sequences containing varying levels of amino acid conservation. Residue-residue couplings drive the coevolution of structurally and functionally connected positions, leading to correlated columns in multiple sequence alignments of homologous proteins. These correlations can be modelled using a Potts model energy framework defined by local biases (h_i_) and pairwise coupling terms (J_ij_). **(b)** Based on the Potts model, a direct information network is constructed to identify nodes with high betweenness centrality, corresponding to key residues in the evolving functional network. Nodes (amino acid positions) are colored by subdomain localization, and edge thickness reflects the strength of direct information (DI). Under low selection pressure (up to round 6), strong couplings are predominantly confined to the fingers domain; as selection pressure increases, stronger couplings emerge in the thumb domain, indicating a shift in coupling pathways during adaptation.

Only a small fraction of these couplings corresponds to classic structural contacts, defined here as residue pairs with C_α_-C_α_ distances below 8 Å, so they are better described as functional couplings **(Supplementary Figure 8)**. Repeatedly, positions 607 in Motif A and 749 in Motif B served as coupling hubs. They were consistently implicated at every evolutionary stage despite changing partner positions. Some strengthened couplings, such as between 732, 739, and 749, emerged after four directed evolution rounds from IE004 to IE010. The progressive enhanced selective pressure led to new couplings, such as 607 and 655, or losses of others because specific sites reached almost 100% mutational rate (such as the 751-753 coupling). New functional coupling patterns involving positions in the palm and thumb domains appeared later in directed evolution cycles and replaced couplings in the fingers subdomain. For instance, strong connections appeared between residues 505 and 554 or 505 and 518 in IE013. The top mutational hotspots (L505, L554, A607, I616, V739, M749, F751, M753) also ranked among the most functionally coupled positions, spotlighting critical residues supporting new polymerase activity. Epistatic coupling analysis thus enables precise identification of key mutational networks switching progressively from fingers to thumb domains and supporting polymerase evolution. Among the strongest couplings, which are also observed in variant E5, are the residue pairs 751-753, 739-749, 616-607, and 554-505 (**Figure 7b**).

To better characterize the outcome of the directed evolution of the Tth DNA polymerase, a quantitative fitness score was assigned to each variant to generate sequence-function pairs. Fitness values were estimated by comparing the frequency of each variant in the selected library to its frequency in the initial unselected library and were normalized to the enrichment of the WT polymerase. To test whether mutations accumulated during evolution act approximately additively on an unobserved latent trait, a global epistasis model, following the work of Otwinowski et al., was fitted to the sequence-function data^45^. The model provided an excellent fit to the variant sequences and their measured fitness in the final selected library (65,513 variants; *r*^2^ = 0.87, *σ*^2^ = 0.15) **(Figure 8a**, **Figure 8b)**. The epistatic model fitted the experimental fitness with a slightly higher r-squared than for a purely additive (non-epistatic) model (*r*^2^ = 0.87 versus *r*^2^ = 0.83, *σ*^2^ = 0.20), indicating that incorporating a nonlinear mapping from the latent additive trait to observed fitness captures additional structure in the genotype-fitness landscape.

**Figure 8.**
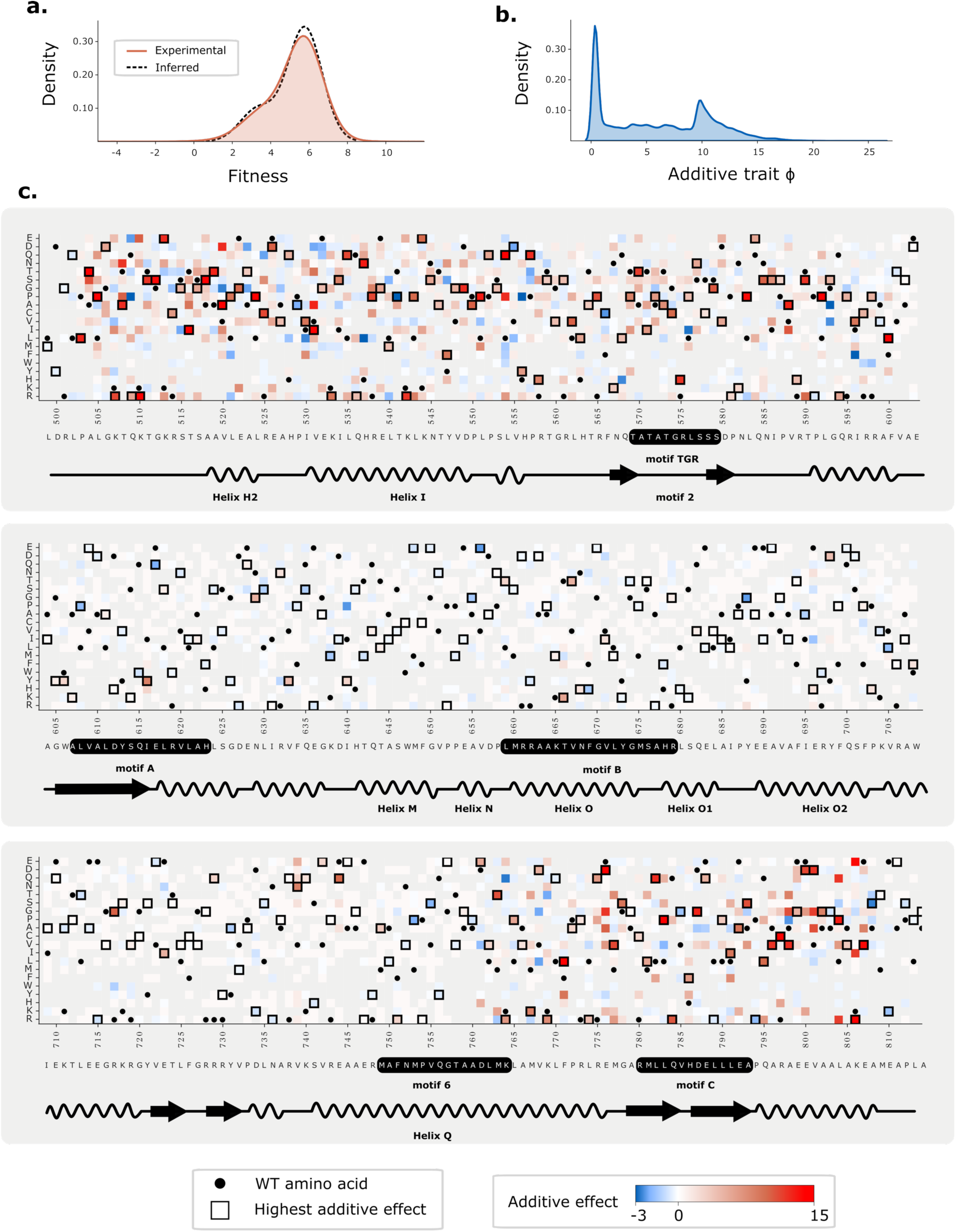
Global epistasis inferred with a flexible monotonic I-spline model. The dataset corresponding to the final selected library (65,513 variants) was analyzed using the global epistasis framework of Otwinowski et al. ^45^, in which the observed fitness is modeled as a nonlinear monotonic function *g*(*φ*) of an underlying additive trait *φ* defined as a sum of third-order I-splines. The additive trait *φ* is defined as the sum of positive or negative additive contributions of each mutation, and both the I-spline coefficients and additive effects are inferred jointly by maximum likelihood. The resulting model explains very well the experimental fitness (r^2^ = 0.87, σ^2^= 0.15). **(a)** Distribution of fitness values, quantified as the log enrichment ratio of each variant relative to the WT, showing a bimodal pattern. The solid line denotes the experimentally measured distribution and the dashed line the distribution predicted by the inferred global epistasis model. **(b)** Distribution of the additive trait *φ*, which is strongly multimodal with predominantly positive values, consistent with most variants having high fitness in the final round of evolution. For visualization, *φ* is rescaled so that the WT trait is zero, following the convention of Otwinowski et al.^45^. **(c)** Heat map of the inferred additive effect of each amino-acid substitution on the underlying trait *φ*. Black circles mark WT residues at each position, and squares indicate the substitution with the maximal absolute additive effect at that position. Additive effects are largely concentrated in the finger region (positions < 600) and in the palm region (positions > 770), with comparatively few large-effect substitutions in other domains.

Among the 10 mutations with the largest positive effects on the inferred additive trait, two correspond to substitutions present in variant E5 (L505P and L554Q), and none of the E5 mutations were inferred to have a negative effect in this model. The positions with the strongest epistatic effects are located in the thumb and palm domains, as identified by the most coupled residues in the mfDCA coevolution analysis of the final library. **(Figure 8c)**.

Overall, these results indicate that the sequence-function relationships can be described adequately by a global epistasis model and direct coupling analysis (DCA). Most variations are explained by additive contributions of individual mutations modulated by a nonlinear mapping. However, it appears that the epistasis model is not exhaustive, and a residual fraction of unexplained variance likely reflects higher-order or long-range couplings between mutations that are not captured by the global epistasis approximation.

### Fitness landscape and epistatic clusters

To fully characterize the directed evolution on Tth polymerase, a fitness landscape was generated from sequence:function pairs. A total of 194,000 unique variants with a fitness score were projected on a two-dimensional space, enabling detection of higher-order mutation couplings. Successive libraries displayed increasing fitness histograms, indicating progressive enrichment in active variants. Sequences were embedded using ESM2-3B^46^, visualized in two dimensions, and interpreted via fitness scores. This allowed a mapping of mutation effects at the level of single amino acid resolution (**Figure 9a**).

**Figure 9.**
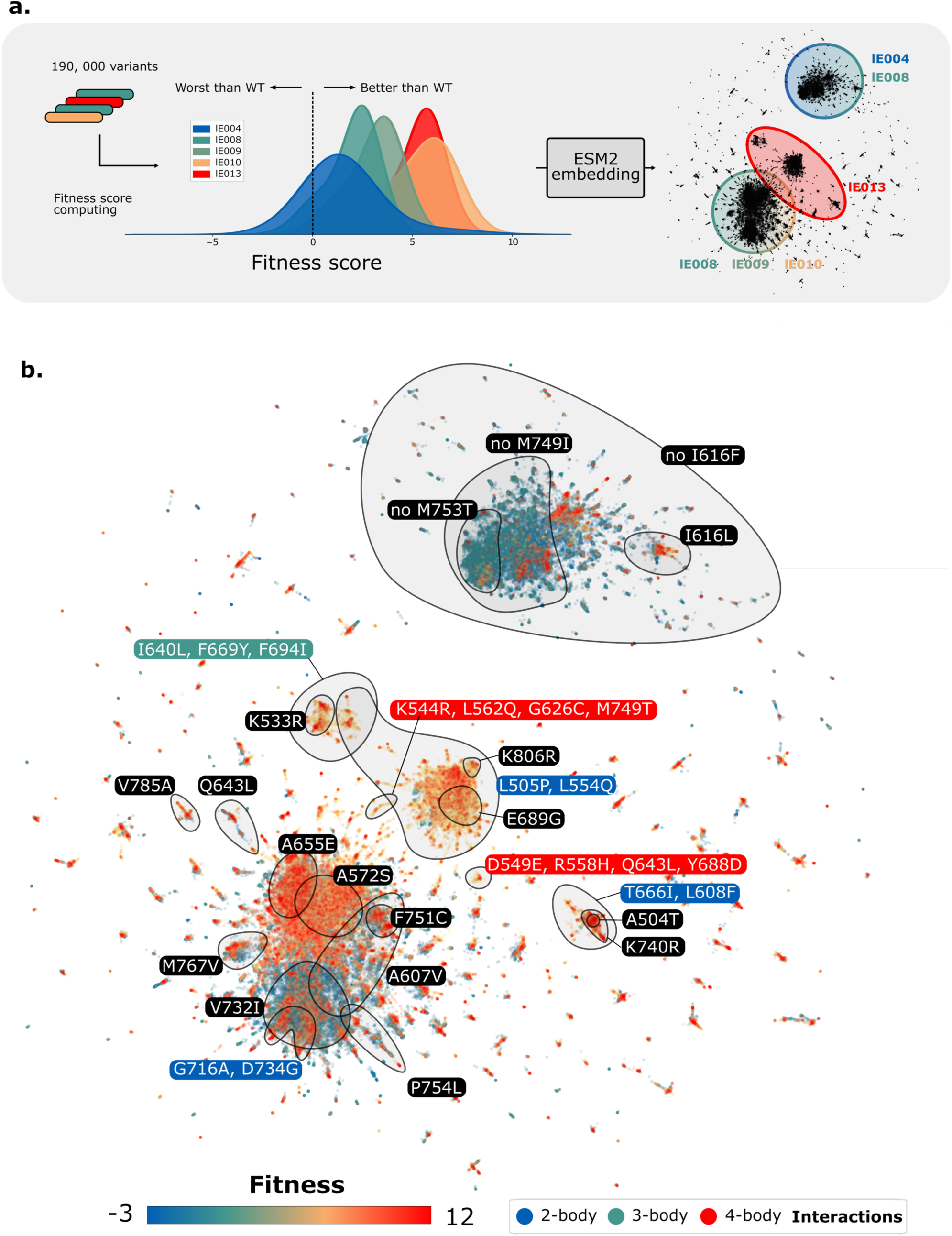
Evolutionary fitness landscape inferred from individual variant frequencies and protein language model embedding. **(a)** Nanopore sequencing enables tracking of variant frequencies for approximately 194,000 polymerase variants across the course of evolution, allowing estimation of individual fitness scores. Dimensionality reduction using the ESM-2 (6B) protein language model and a UMAP projection provides a latent fitness landscape representation. The most populated regions of sequence space shift progressively between libraries while retaining substantial overlap, reflecting continuous evolutionary trajectories with a progressive sequence space exploration. **(b)** The ESM-2 sequence embedding enables residue-level visualization of the fitness landscape. Among the 100 most frequent mutations in the final library, clusters of co-occurring variants can be identified. This mapping reveals higher-order couplings beyond pairwise interactions, including 2-body (blue), 3-body (green; e.g., coordinated variation at positions 640, 669, 694), and 4-body couplings (red; e.g., 544, 562, 626, 749 or 549, 558, 643, 688).

This analysis revealed valleys and peaks of fitness in a particularly rugged landscape, also observed for other enzymes such as yeast dihydrofolate reductase^47^. Variants from the same library consistently clustered within defined zones; libraries 4 and part of 8 formed one cluster, while the remainder of 8, plus libraries 9 and 10, grouped into another, and library 13 formed a distinct cluster. However, there was a substantial overlap between the libraries.

Fitness mapping also revealed evolutionary trajectories as in a phylogenetic tree. The fitness landscape appears mainly as valleys separated by high-energy barriers. The walk toward local fitness peaks were made possible by each error-prone PCR round introducing hundreds of thousands of jumps in all directions of the sequence space. This random exploration of the sequence space through error-prone PCR clearly shows the concentration of polymerase variants with higher fitness in some regions compared to others.

This visualization made it possible to identify important high-order mutation clusters beyond pairwise interactions, which consistently shared signature substitutions. For example, among the hundred most frequent mutations, mutations I640L, F669Y, and F694I co-occurred as a group (three-body cluster), as did D549E, R558H, Q643L, and Y688D, as well as K544R, L562Q, G626C and M749T, highlighting the existence of 4-body functional couplings (**Figure 9b**). Most of these mutations are conservative. However, their positions are separated by large distances that span both the fingers and thumb domains of the polymerase. The cluster comprising residues 549, 558, 643, and 688 is illustrative: positions 643 and 688 lie at the tip of the fingers domain, whereas 549 and 558 are located at the base of the thumb domain, with these two regions separated by a maximum of 50 Å. Similarly, the two residue pairs forming the second cluster (544, 562 and 626, 749) are separated by 45 Å maximum. In that sense, this method goes beyond the Potts model which includes only single-site and pairwise interactions and allows the discovery of long-range couplings **(Supplementary Figure 10)**.

These findings underscore the utility of the improved RT-CSR platform and of next-generation sequencing for trying to understand the underlying evolutionary mechanisms.

## Discussion

To synthesize enzymatically DNA from RNA or XNA, wild-type reverse transcriptases (RTs) from retroviruses are the natural choice but are often suboptimal for many biotechnological applications. RNA-dependent RT function has been improved for fidelity^10,12^ or for kinetics^9^ by screening methods and directed evolution from family-A and family-B DNA polymerases. However, discovering enzymes that copy XNAs remains difficult. Here we show that a variant of RT-CSR can be implemented in a microfluidic one-pot format and coupled with next-generation sequencing to explore evolutionary trajectories. We obtain the first reverse-transcriptase derived from family-A polymerases for copying XNA and notably the bulky 2’MOE modifications over a large range of temperatures.

The one-pot microfluidic RT-CSR provides highly reproducible reaction conditions and simplifies the workflow by eliminating two successive PCR steps. The approach is inherently versatile, because changing only the selection primer to incorporate alternative modified bases allows direct targeting of new substrates. In this work, the evolutionary gap to overcome was particularly large. The WT polymerase stalls for more than an hour in the presence of a single 2’MOE modification and is not able to accommodate properly the modified duplex to pursue reverse-transcription **(Figure 5)**. To minimize biases, we designed a selection oligonucleotide that contains both canonical RNA bases and 2ʹMOE modifications and includes both purines and pyrimidines. The resulting E5 mutant has gained a completely new activity and is now able to bypass this modification. The activity was also improved by a factor 80 for 2’OMe modification **(Figure 3)**. For comparison, in a previous evolution campaign, described by Houlihan et al.^10^, the starting point was an already engineered PolB capable of extending 2’OMe-containing templates by roughly 40 nucleotides. A plausible explanation for the greater engineering difficulty of PolA compared with PolB polymerases is the smaller, more constrained catalytic channel in PolA enzymes, which may limit their capacity to accommodate new substrates as shown in^48,49^. Differences in intrinsic thermostability between thermophilic (PolA) and hyperthermophilic (PolB) polymerases may further restrict the accessible mutational space and thereby exacerbate this contrast in evolvability.

By propagating entire libraries rather than isolating single clones at each cycle, our approach enables a detailed, stepwise analysis of epistasis during evolution while allowing broader (though less exhaustive) exploration of sequence space. The multiplicity of starting points at each diversification cycle facilitates sampling of larger regions in sequence space in search of local fitness peaks and the rescue of possibly interesting variants. This is illustrated by the fluctuation in mutational rates along the evolution **(Figure 6)**. However, given the finite size of the library, this comes at the cost of an increased sparsity. This strategy revealed for the first time local pairwise patterns **(Figure 7)** as well as higher-order interactions spanning up to four residues **(Figure 9)**. Several substitutions are consistent with earlier studies, for example at position 616, which is also mutated in Raghunathan et al.^50^, and at positions 616 and 749 in Chen et al.^8^, and 749 in Vichier-Guerre et al.^9^. Therefore positions 616 and 749 likely play an important role in controlling reverse-transcription activity on these challenging substrates. It is probably even more the case for M749 which we find to be involved in epistasis along evolution cycles and close to the RNA template. Except for these positions, most of the sites uncovered here appear novel and unique. It is not unexpected that 8 of the 13 mutations of E5 involve the 8 positions with the highest frequency in the last database as the final bank is dominated by these mutations (>20%) **(Supplementary Figure 7).** In comparison, the positions involved in the evolution of *polB* for reverse transcription are also mostly unique but share some mutational hotspots. The motif B seems to play a critical role: in Houlihan et al.^10^, mutations are predominantly located within this motif for C8 polymerase. For the RTX polymerase described by Ellefson et al.^12^, the mutations are distributed all along the sequence, but motif B mutations appeared to be strongly enriched along the evolution.

Efficient RTs for C2’-modified nucleotides are of particular interest for sequencing endogenously modified cellular RNAs such as tRNAs or ribosomal RNA that can bear C2’ methylation after transcription^51^. RT-based technologies such as Illumina fail when standard enzymes cannot process modified templates, which then remain invisible during the RNA-seq analysis. The identified E5 polymerase should also prove valuable for SELEX and aptamers selection. Although it does not support fully 2ʹMOE-modified templates and could require further engineering, it efficiently reverse transcribes partially modified libraries. It enables most applications that require only limited 2ʹMOE or 2’OMe reverse transcription such as the final reverse transcription step required before DNA amplification in SELEX selection^5,6^. E5 polymerase can also support the high demand of reverse-transcriptases in combination with Cas9 nickase for genome editing such as in “prime editing”^52^.

Some polymerases from the A family already possess reverse transcriptase activity but are not as thermostable, efficient and versatile as the variant discovered in this study, which greatly limits their use and storage^53,54^. Overall, this study delivers a PolA reverse transcriptase fully active on RNA, also active on 2’OMe and 2’MOE-modified substrates and with an improved thermal stability.

The method generated one of the largest evolutionary datasets available for a DNA polymerase, encompassing more than 194,000 variants associated with a frequency-based fitness score **(Supplementary Table 3)**. In the near future, a more advanced analysis of these original libraries will likely become possible as more computing techniques are developed using deep learning models, and this may be useful for training them in the in silico design of smarter libraries for enzymes dealing with other XNA substrates^55^. To go beyond the E5 variant, a rational approach could leverage the eight most frequent mutations from the library as a common basis, augmented by those with the strongest positive epistatic effects on the trait, as identified in the analysis inspired from Otwinowski et al.^45^. Alternatively, to select a better variant in the final library, large language models could provide embeddings to train a discriminator on a smaller set of known variants with experimental characterization, followed by scoring of the entire library aligned with the highly effective technique demonstrated in EVOLVEpro^56^.

## Methods

### Design and construction of mutagenic libraries

Mutagenic error-prone PCRs (epPCR) were performed using Hot Start Taq DNA polymerase (NEB M0495). The protocol was established following Martineau et al.^57^, with biased nucleotides concentrations (350 μM dATP, 400 μM dCTP, 200 μM dGTP, and 1350 μM dTTP), 0.5 mM MnCl₂, 0.1 nM template, 200 nM forward primer, 200 nM reverse primer, and 1.6% Hot Start Taq polymerase mixed in the reaction buffer (10 mM Tris-HCl, pH 8.3, 50 mM KCl, 2.5 mM MgCl_2_). PCR cycling conditions were as follows: initial denaturation at 95 °C for 2 minutes, 10 cycles of denaturation at 95 °C for 30 seconds, annealing at 55 °C for 30 seconds and extension at 68 °C for 3 minutes. The products were subsequently analyzed on agarose gels. The products were incubated with rCutsmart buffer 1x (NEB B6004) and DpnI endonuclease (NEB R0176S) for 60 minutes at 37 °C and 20 minutes at 80 °C to remove methylated parental DNA.

A second PCR was performed to include overlap regions using Q5 polymerase with Q5 PCR buffer 1X (NEB B9027), GC Enhancer 1X (NEB B9028), 200 μM dNTPs (NEB N0447), 200 nM forward primer, 200 nM reverse primer, 2 μL of epPCR product, and 4 units Q5 (NEB M0491S). PCR cycling conditions were as follows: initial denaturation at 98 °C for 2 minutes, 23 cycles of denaturation at 95 °C for 10 seconds, annealing at 68 °C for 30 seconds and extension at 72 °C for 1 minute. The product was purified on an agarose gel and then using the MinElute kit (Qiagen 28004).

Cloning was proceeded by Gibson assembly with the NEBuilder HiFi kit (NEB E2621), using 90 fmol insert and 30 fmol backbone, incubated at 50 °C for 60 minutes. Three microliters of the product were transformed into 100 μL of NEB10beta electrocompetent cells (NEB C3020). The cells were grown at 37 °C overnight into 100 mL LB-kanamycin medium (50 µg.mL^-1^). The sizes of the libraries were assessed by spreading serial dilutions on LB-kanamycin-agar plates.

All oligonucleotides used in this study are detailed in **Supplementary Table 2**.

### One-pot RT-CSR selection

The library cloned in pET-29a(+) (Novagen) was expressed in BL21 star^TM^ (DE3) (Invitrogen^TM^ C601003) cells induced for 3 hours at 37 °C after they reached OD_600_ 0.6. The pellet, stored at -20 °C, was thawed and resuspended in 500 μL Tth buffer 1x, defined as 10 mM Tris-HCl pH 8.4, 50 mM KCl, 1.5 mM MgCl₂, 0.1% Triton. Cells were pelleted at 3,500 × g for 5 minutes and washed again. The reaction mix used NEB standard Taq buffer (NEB B9014) supplemented with 0.1% Triton X-100, 200 μM dNTPs, 40 nM selection forward and reverse primers, 200 nM outnested biotinylated forward and reverse primers, 10 ng.μL^-1^ yeast RNA (Roche), 0.4% Pluronic F-127 (Invitrogen), 0.2 mg.mL^-1^ BSA (Sigma-Aldrich), and washed cells at final OD 0.025. Water-in-oil droplets emulsions were generated using a microfluidic flow-focusing design with Novec HFE 7500 fluorinated oil (3M^TM^) supplemented with 4% Fluosurf-C^TM^ surfactant (Emulseo) as continuous phase. The microfluidic masks were designed on LibreCAD and ordered from Selba. The resulting two-step mold was realized by soft lithography following SU8 protocols (Microchem) and using an MJB4 mask aligner (SUSS) in a clean room. The microfluidic chips were obtained by bonding the inlet-punched imprinted PDMS slabs on microscopy glass slides after O_2_ plasma functionalization. Channel hydrophobicity was ensured by pre-treating chips with Novec HFE 1720 (3M), followed by incubation at 110 °C for 10 minutes and cooling at room temperature.

Emulsions were collected and subjected to PCR cycles with 5 minutes initial denaturation at 94 °C, then 27 cycles 30 seconds denaturation at 95 °C, 30 seconds annealing at 60 °C and 3 minutes extension at 72 °C. To recover PCR products, PCR tubes were washed twice with 100 μL Novec HFE 7500 (3M), vortexed, and centrifuged (7,500 × g, 2 min). Bottom oil phase was then removed. Next, 100 μL water and 200 μL Perfluorooctanol (Sigma-Aldrich) were added, vortexed, and centrifuged. The top aqueous supernatant was finally collected.

Biotin-streptavidin separation was performed incubating 5 μL Dynabeads C1 (Invitrogen^TM^) for 1 hour in buffer (20 mM Tris-HCl pH 7.5, 2 M NaCl, 0.2% Tween 20, 2 mM EDTA) after two washing steps with 500 μL of this buffer. Beads were concentrated in 50 μL, then mixed with emulsion PCR aqueous phase. After 1-hour incubation, beads were washed twice with 200 μL 30 mM NaOH (37 °C), once with 200 μL (10 mM Tris-HCl pH 8.8, 0.01% Tween 20, 0.1 mM EDTA), finally resuspended in 10 μL 10 mM Tris-HCl pH 8.8. A Q5 PCR as described in the previous section was used to add overlap regions required for Gibson assembly.

### Polymerase activity screening

Libraries were transformed into BL21 star^TM^ (DE3) *E. coli* and plated onto kanamycin LB agar plate (50 µg.mL^-1^). Single colonies were transferred into 96-well plates (Eppendorf) containing 1 mL LB-kanamycin medium and cultivated overnight at 37 °C with shaking at 800 rpm using an Eppendorf ThermoMixer incubator. A 10 μL aliquot of overnight culture was inoculated into fresh 1 mL kanamycin LB medium (50 µg.mL^-1^), and cells were grown to OD_600_≈0.6. Expression was induced with 1 mM IPTG, and cultures were incubated at 20 °C, 800 rpm for 12 h. Cells were pelleted by centrifugation (3,200 × g, 25 °C, 20 min), resuspended in 50 μL 1× Tth buffer (10 mM Tris-HCl pH 8.4, 50 mM KCl, 1.5 mM MgCl₂, 0.1% Triton X-100), and transferred to 96-well PCR plates. Samples were incubated at 75°C in an Eppendorf ThermoMixer for 30 min, cooled on ice for 15 min, and centrifuged (3,200 × g, 25 °C, 1 h). To assess polymerase activity, reactions were assembled to a final volume of 10 μL containing 1× Standard Taq buffer, 1 μM modified template, 1 μM 5’-FAM-labelled primer, 1 mM dNTPs, and 1 μL lysate, and incubated at 72 °C for 1 h. Reactions were terminated by the addition of two volumes of stop buffer (10 mM EDTA, 98% formamide, 1 mg.mL^-1^ bromophenol blue, 10 μM unlabeled complementary primer). Prior to analysis, samples were preheated at 95°C for 10 min, resolved by denaturing 18% urea-polyacrylamide gel electrophoresis, and visualized using a Typhoon FLA 9000 biomolecular imager (Cytiva).

### Polymerases purification

BL21 star^TM^ (DE3) competent cells were transformed with the pET-29a(+) Tth expression plasmid. A selected colony was inoculated into LB medium and cultured at 37 °C. Protein expression was induced by adding 1 mM IPTG when the culture reached OD_600_ 0.6, followed by an overnight incubation of the 1 L culture at 20 °C. Cells were harvested by centrifugation at 3,300 × g for 30 min, and the pellet was resuspended in 40 mL lysis buffer (50 mM Tris-HCl pH 7.5, 300 mM NaCl, 1 mM DTT, 0.05% Triton X-100). One tablet of protease inhibitor cocktail (Thermo Scientific) and 1 μL benzonase nuclease (Millipore) per 30 mL of lysate were added. Cell disruption was performed by sonication with five cycles (60s on - 60s off). Lysates were clarified at 30,000 × g for 30 min.

Polyethylene glycol (PEG) 8000 was added to the supernatant to a final concentration of 10%. The solution was heated at 70 °C for 15 min, cooled for 5 min at room temperature and on ice for 15 min. A white/yellow precipitate formed in the lysate, which was removed by centrifugation at 15,000 × g for 40 min. The supernatant was collected and diluted to adjust the salt concentration to approximately 90 mM before HiTrap Heparin purification (Cytiva). The Heparin elution was performed using a gradient between buffer A (100 mM NaCl) and buffer B (1 M NaCl). Eluted fractions were dialyzed in storage buffer (10 mM Tris-HCl pH 7.5, 100 mM NaCl, 1 mM DTT, 0.1 mM EDTA, 10% glycerol).

Protein purity was analyzed by SDS-PAGE **(Supplementary Figure 3)** using 4-15% or 4-12% gels with a Precision Plus Protein molecular weight ladder (BioRad). Enzyme samples were concentrated with Amicon Ultra centrifugal filters (30 kDa MWCO, Merck), flash frozen in liquid nitrogen, and stored at -20 °C.

### Polymerases kinetics

E5 and WT polymerases were purified as described above. To quantify their kinetics across the modified bases, primer-extension reactions were analyzed by capillary electrophoresis. Reactions (10 µL) contained 1× Standard Taq buffer, 0.5 µM 5ʹ-ATTO488-labelled primer annealed to the modified template, 1 mM dNTPs, and 1 µM purified polymerase (E5 or WT), and were incubated at 72 °C for 1, 2, 5, 10, 30 and 60 min. Each time course was performed in duplicate. Reactions were quenched by adding 40 µL 50 mM EDTA, diluted 1:100 in milliQ water. Samples were submitted to Microsynth for fragment-length analysis by capillary electrophoresis (Filter set : DS-33_G5, Size standard : Genescan 120 LIZ). Traces were aligned and peak areas were used to calculate the fraction of fully extended primer relative to partially extended and unreacted species, yielding the extent of product formation at each time point.

### Libraries tagging and sequencing

Oxford Nanopore Technology (ONT) sequencing is limited by its high error rate, which precludes confident single nucleotide polymorphism (SNP) detection. The approach proposed by Karst et al.^58^ addresses this limitation through the introduction of paired unique molecular identifiers (UMIs) to generate high-accuracy consensus sequences. We adapted their protocol and extended the bioinformatic analysis pipeline to suit our specific experimental requirements. As described in **Supplementary Figure 9a**, a Q5 PCR was performed to introduce 44-nt semi-structured UMI flanking the region of interest such that all reads sharing the same pair of UMI derive from a single input molecule. PCR products were recloned using Gibson assembly as described before. The resulting library was transformed into NEB 10-beta electrocompetent cells (C3020) cells by electroporation and transferred into 100 mL LB medium supplemented with kanamycin (50 µg.mL^-1^) for overnight growth at 37 °C. Following library amplification, PCR was performed to generate sequencing-ready DNA using the following conditions: Q5 PCR buffer 1x, (NEB B9027), GC Enhancer 1x (NEB B9028), 200 μM dNTPs (N0447), 200 nM forward primer, 200 nM reverse primer, 1-5 ng barcoded plasmid DNA, and 10 units Q5 polymerase (M0491S). Cycling conditions were as follows: initial denaturation at 98 °C for 2 min; 27 cycles of 98 °C for 10 s, 68 °C for 30 s, and 72 °C for 1.5 min. 500 μL reaction mix was made and column purified using the MinElute kit (Qiagen 28004). A low cycle number combined with prolonged extension times was used to minimize PCR-mediated chimera formation between variants, which would otherwise distort quantitative analysis of variant representation in the library.

Sequencing libraries were prepared using the Ligation Sequencing Amplicons V14 protocol (SQK-LSK114). Sequencing was stopped once the sequencing read count reached 50-fold sample coverage (assuming 10^5^ variants and 2 kb amplicon lengths) corresponding to >10 Gb sequencing data and around 5 million reads.

### Nanopore sequencing denoising

The Nanopore sequencing denoising workflow is shown in **Supplementary Figure 9**. The pipeline adapts previously described long-read denoising strategies^58–61^ to much larger datasets **(Supplementary Table 3)**. Briefly, raw ONT signals are basecalled with Dorado (SUP v5.0.0), and reads are aligned to the Tth reference fragment using Minimap2^62^. Reads are filtered by length and mapping quality. UMI sequence pairs of each read are extracted from the alignments and clustered with CD-HIT-EST^63^ at 91.5% identity, thereby grouping reads derived from the same molecule. Clusters supported by fewer than five reads are discarded. Within each remaining cluster, variants relative to the reference are called with FreeBayes^64^. Mutations are restricted to calls that pass filters on allele fraction while sites outside the mutagenized region of interest and indels are excluded from the analysis. The resulting high-confidence mutations are expected with at least a Phred quality score of 48.7 estimated with a robust benchmarking protocol. Details can be found in supplementary documents.

The sequencing experiments are summarized in **Supplementary Table 3**. The table reports 44 millions of raw reads before clustering and 28 million aligned to the reference. 421,000 clusters (defined as UMI pairs with ≥5 supporting reads) are distributed across 196,000 unique variants recovered with no premature stop codons. Each unique construct is typically covered by an average of 66 reads (mean cluster size across libraries and median=26), providing multiple independent observations of each sequence and enabling robust consensus calling.

### Protein crystallization and structure determination

A truncated version of WT Tth polymerase was generated using the KLD enzyme mix (M0554) after PCR amplification excluding the 5’-3’ exonuclease domain, resulting in only Klenow fragment cloned in pET-29a(+). The enzyme was purified as previously described. Heparin chromatography was followed by size exclusion chromatography on a GF S75 column using buffer containing 50 mM Tris-HCl pH 8 and 150 mM NaCl. The modified DNA substrate was prepared by annealing an 11-mer modified template oligonucleotide with a 16-mer upstream primer oligonucleotide as in^15^. The protein:duplex pair (at a ratio of 1:1.1) was then incubated with 1 mM ddCTP and 10 mM MgCl_2_ in the GF buffer for 30 minutes at room temperature.

Crystallization drops were set up at 18 °C by mixing in a 1:1 ratio the Tth:DNA:dNTP ternary complexes with a crystallization solution consisting of 20% w/v PEG 8000, 0.1 M CAPS pH 10.5 and 0.2 M NaCl. The resulting crystals were flash-frozen in liquid nitrogen after being cryoprotected with the crystallization solution supplemented with glycerol. X-ray crystallographic data were collected at the SOLEIL synchrotron (Saint-Aubin, France) on beamline PX2A. The data were processed using the autoPROC pipeline including Staraniso^65,66^. Subsequently, the structures were solved by Molrep^67^ molecular replacement using an Alphafold^68^ AF2 model as template and refined in Phenix^69^, with manual reconstruction performed using Coot^70^. Details for the dataset are summarized in **Supplementary Table 1.**

### Sequence data analysis

Protein sequences were first filtered to remove all sequences containing an internal stop codon, which are most likely inactive. For each alignment position *i*, the Shannon entropy was computed from the empirical amino-acid frequency distribution *p_i_*(*a*).

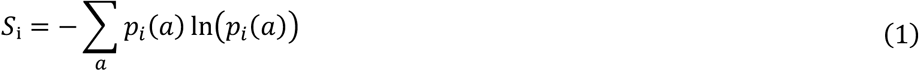

The resulting multiple sequence alignments were already of high quality and were therefore used directly for mean-field Direct Coupling Analysis (mfDCA). Calculations were performed in Python using the py-mfdca implementation of the Matlab method published by Morcos et al.^44^ with default parameters. The DCA model assigns a probability to a sequence of length L based on the observed single-site *a_i_* and pairwise \**a_i_*, *a*_j_+statistics of amino acids across positions.

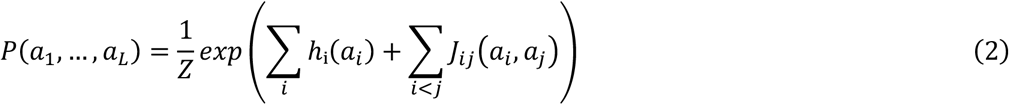

From the inferred model parameters, the direct information between residue pairs was computed to quantify direct coevolutionary couplings with the same py-mfdca package.

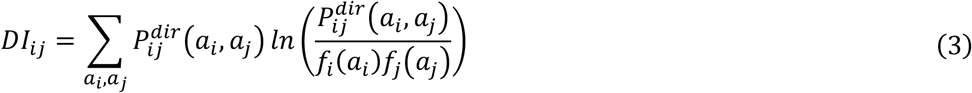

Finally, networks were constructed from these couplings (python package : networkx), and nodes were ranked by their betweenness centrality, defined as the fraction of all shortest paths between any pair of nodes that pass through a given node. Nodes with the highest betweenness act as major intermediates to connect amino acids clusters and were identified as key communication hubs (**Figure 7**).

Sequence counts were obtained for each variant across the libraries, and a frequency was assigned to each variant at its final appearance in the evolutionary trajectory. Enrichment was then calculated for each variant relative to the initial, non-selected library lE001. This enrichment value was subsequently normalized by the enrichment of the WT to obtain a fitness score, which was expressed as the log2 of this normalized enrichment, as previously described^71^. Values below 0 indicate variants with lower fitness than the WT (**Figure 8**, **Figure 9**).

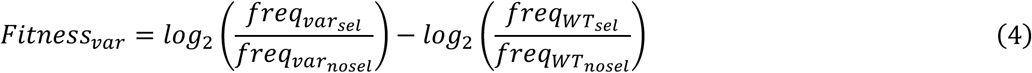

From these experimental fitness values, an explicit statistical model was fitted to estimate the additive contribution of each mutation following the global epistasis framework of Otwinowski et al.^45^. The original implementation was used with hyperparameters *nk* = 4 (number of I-spline knots) and *tol* = 1 × 10^-8^ (convergence tolerance). This model assumes that mutations act additively on an unobserved latent trait ϕ, while the measured fitness is obtained through a monotonic non-linear mapping g(ϕ) represented by the sum of third-order I-splines. The final evolved library, comprising 65,513 unique sequence variants, was used as input. A purely additive (non-epistatic) model was first fitted to the data, yielding a coefficient of determination of *r*^2^ = 0.83. Including a global epistasis non-linearity *g*(*φ*) led to an improved fit with *r*^2^ = 0.87, indicating a modest but consistent gain in explanatory power when epistatic effects are taken into account, in line with previous applications of this framework. The final global-epistasis model accounted for a large fraction of the observed fitness, with residual variance σ^2^=0.15 (versus 0.20 for the non-epistatic model).These metrics indicate that the inferred latent additive effects and the non-linear mapping *g*(*φ*) capture most of the structure in the genotype-fitness landscape while leaving only a limited amount of unexplained higher-order or idiosyncratic epistasis (**Figure 8**).

The variable regions of the sequences were then isolated to accelerate processing and encoded with the esm.pretrained.esm2_t48_3B_UR50D() protein language model^46^. For the analysis, representations from the 30^th^ encoder layer were extracted. After extracting each variant’s embedding, these were reduced in dimensionality through UMAP (python package: sklearn, default settings, n_components = 2). In **Figure 9b** each variant was then colored according to the magnitude of its calculated fitness inspired from^72^.

## Data Availability

*The diffraction data and crystal structure have been deposited in the PDB under the code 30GW. The sequence datasets will be made available from the authors upon reasonable request*.

## Code Availability

Unique Molecular Identifier pairs were extracted from filtered alignments using our in-house tool xumi v1.0.3 deposited in https://github.com/Fravadona/xumi.

## Acknowledgements

The authors acknowledge enlightening discussions with all past and present members of the Delarue and Sauguet teams. They acknowledge the PFBMI facility, the UTechS Single Cell Biomarkers, the Crystallography and the Biomics platforms, the Bioinformatics HUB at Institut Pasteur for their technical support and advices. The authors thank the Biomaterials & Microfluidic core facility (BMcf) at Institut Pasteur for their support with the production of the microfluidic chips. They acknowledge SOLEIL for the provision of synchrotron radiation facilities and the staff of beamlines PROXIMA-1 and PROXIMA-2 (Saint-Aubin, France). This work was supported by the MoleculArXiv PEPR and PICNA-APT (ANR-23-CE44-0033) ANR grants. The ADMB unit is an *Équipe labellisée* of the Fondation pour la Recherche Médicale (FRM). EV was funded by Université Paris Cité.

## Ethics declarations

Competing interests: None declared.

## Contributions

M.D., E.V. and R.S. conceived the project and designed the experiments. E.V. evolved the enzyme. E.V. and C.L. performed the biochemical characterization. E.V. and A.H. contributed to protein structure determination. E.V, R.N., R.S. and I.C. performed sequencing experiments and analysis. E.V. and M.D wrote the paper. All authors reviewed and commented on the paper.

## Notes

### Competing Interest Statement

The authors have declared no competing interest.

