## Supplementary Data and Figures for "Directed evolution of a thermostable family-A DNA polymerase into a reverse transcriptase for 2’-modified RNA"

#### Supplementary Information

Eloi Vincent[1], Clément Lopez[1], Rémi Sieskind[1], Isciane Commenge[1], Rafael Navaza[2], Ahmed Haouz[2], Marc Delarue[1]

1. Institut Pasteur, Université Paris Cité, CNRS UMR 3528, Unit of Architecture and Dynamics of Biological Macromolecules, 25 rue du Docteur Roux, 75015 Paris, France

2. Institut Pasteur, Université Paris Cité, CNRS UMR 3528, Plate-forme de Cristallographie-C2RT, 28 rue du Docteur Roux, 75724 Paris Cedex 15, France

### Supplementary data

#### Protein sequences

##### WT protein sequence

MEAMLPLFEPKGRVLLVDGHHLAYRTFFALKGLTTSRGEPVQAVYGFAKSLLKALKEDGYKAVFVVFDAKAPSFRHE
AYEAYKAGRAPTPEDFPRQLALIKELVDLLGFTRLEVPGYEADDVLATLAKKAEKEGYEVRILTADRDLYQLVSDRV
AVLHPEGHLITPEWLWEKYGLRPEQWVDFRALVGDPSDNLPGVKGIGEKTALKLLKEWGSLENLLKNLDRVKPENVR
EKIKAHLEDLRLSLELSRVRTDLPLEVDLAQGREPDREGLRAFLERLEFGSLLHEFGLLEAPAPLEEAPWPPPEGAF
VGFVLSRPEPMWAELKALAACRDGRVHRAADPLAGLKDLKEVRGLLAKDLAVLASREGLDLVPGDDPMLLAYLLDPS
NTTPEGVARRYGGEWTEDAAHRALLSERLHRNLLKRLEGEEKLLWLYHEVEKPLSRVLAHMEATGVRLDVAYLQALS
LELAEEIRRLEEEVFRLAGHPFNLNSRDQLERVLFDELRLPALGKTQKTGKRSTSAAVLEALREAHPIVEKILQHRE
LTKLKNTYVDPLPSLVHPRTGRLHTRFNQTATATGRLSSSDPNLQNIPVRTPLGQRIRRAFVAEAGWALVALDYSQI
ELRVLAHLSGDENLIRVFQEGKDIHTQTASWMFGVPPEAVDPLMRRAAKTVNFGVLYGMSAHRLSQELAIPYEEAVA
FIERYFQSFPKVRAWIEKTLEEGRKRGYVETLFGRRRYVPDLNARVKSVREAAERMAFNMPVQGTAADLMKLAMVKL
FPRLREMGARMLLQVHDELLLEAPQARAEEVAALAKEAMEKAYPLAVPLEVEVGMGEDWLSAKG

##### E5 protein sequence

MEAMLPLFEPKGRVLLVDGHHLAYRTFFALKGLTTSRGEPVQAVYGFAKSLLKALKEDGYKAVFVVFDAKAPSFRHE
AYEAYKAGRAPTPEDFPRQLALIKELVDLLGFTRLEVPGYEADDVLATLAKKAEKEGYEVRILTADRDLYQLVSDRV
AVLHPEGHLITPEWLWEKYGLRPEQWVDFRALVGDPSDNLPGVKGIGEKTALKLLKEWGSLENLLKNLDRVKPENVR
EKIKAHLEDLRLSLELSRVRTDLPLEVDLAQGREPDREGLRAFLERLEFGSLLHEFGLLEAPAPLEEAPWPPPEGAF
VGFVLSRPEPMWAELKALAACRDGRVHRAADPLAGLKDLKEVRGLLAKDLAVLASREGLDLVPGDDPMLLAYLLDPS
NTTPEGVARRYGGEWTEDAAHRALLSERLHRNLLKRLEGEEKLLWLYHEVEKPLSRVLAHMEATGVRLDVAYLQALS
LELAEEIRRLEEEVFRLAGHPFNLNSRDQLERVLFDELRLPAPGKTQKTGKRSTSAAVLEALREAHPIVEKILQHRE
LTKLKNTYVDPLSSQVHPRTGRLHTRFNQTATATGRLSSSDPNLQNIPVRTPLGQRIRRAFVAEAGWTLVALDYSQF
ELRVLAHLSGDENLIRVFQEGKDIHTQTASWMFGVPPEAVDPLMRRAAKTVNFGVLYGMSAHRLSQELAIPYEEAVA
FIERNFQSFPKVRAWIVKTLEEGRKRGYVETLFGRRRYVPDLNARDKSAREAAERIAYNTPVQGTAADLMKLAMVKL
FPRLREMGARMLLQVHDELLLEAPQERAEEVAALAKEAMEKAYPLAVPLEVEVGMGEDWLSAKG

#### Nanopore sequencing data processing

##### *1. Basecalling and mapping*

Oxford Nanopore Technologies (ONT) raw signal files were basecalled using Dorado v0.9.0 (https://github.com/nanoporetech/dorado; model dna_r10.4.1_e8.2_400bps_sup@v5.0.0) with default parameters. Reads were aligned to the reference construct containing all regions present in the sequenced reads, including a unique molecular identifier (UMI) pair and the mutagenized region of interest (ROI), using minimap2 v2.27-r4, integrated within Dorado[1,2].

##### *2. Read filtering*

Aligned reads were filtered using SAMtools v1.16[3]. Only primary alignments with mapping quality (MAPQ) > 20 and read length within ±25% of the expected construct length, adjusted per dataset, were retained. These criteria remove low-quality mappings and reads that are unusually short or long, including putative concatemer artifacts.

##### 3. UMI extraction

UMI pairs were extracted from filtered alignments using our in-house tool xumi v1.0.3 (https://github.com/Fravadona/xumi, doi:10.5281/zenodo.18905757), which extracts per-read subsequences at user-specified reference regions. Each UMI was required to be 44 ±6 bp. Retained UMI pairs were concatenated and written as FASTA sequences identified by their source read's query name. The latter two steps were performed using a custom AWK script.

##### 4. UMI clustering

Concatenated UMI sequences were clustered using CD-HIT-EST v4.8.1[4] at 91.5% sequence identity in high accuracy mode (-g 0) while preserving full sequence identifiers (-d 0) to allow mapping of clusters back to the original reads in the SAM file. Given the $3^{64} \approx 3.4 \times 10^{30}$ sequence space of the UMI pair design (64 degenerate bases out of 88, each allowing 3 possible nucleotides), the probability of incorrectly merging distinct UMI sequences at this threshold is vanishingly small relative to experimental library sizes ($\approx 10^5$).

##### 5. Cluster filtering

A minimum depth of 15 reads has been recommended for consensus generation from UMI-grouped reads using earlier ONT chemistries[5]. Our benchmarking on R10.4.1 ONT data shows that consensus accuracy in the mutagenized ROI stabilizes at depth ≥ 5 (see "Benchmarking cluster depths"). Only clusters meeting this threshold were retained.

##### 6. Per-cluster contig construction

Using a custom Python script, each retained cluster was first linked to its constituent reads via the CD-HIT-EST cluster file. Matching reads were then extracted from the alignment file and assembled into separate reference contigs within a single SAM file to enable independent variant calling. A companion FASTA file was generated by copying the original reference sequence under each contig’s name. Finally, the SAM file was sorted and indexed using SAMtools.

##### 7. Variant calling

Variants were called from the clustered alignments using FreeBayes v1.3.8[6] with default parameters, except that ploidy was set to 1. FreeBayes was chosen for its direct compatibility with BAM alignments, its haploid mod, which is appropriate since each cluster represents reads from a single UMI-tagged molecule, and its standard VCF file format output.

##### 8. Variant filtering and normalization

The VCF was filtered using BCFtools v1.16[3], retaining only variants where the alternate haplotype observation count (AO) exceeded the reference haplotype observation count (RO) (i.e. AO > RO). At positions with multiple candidate alleles, the best-supported allele was selected based on AO, using QA (the sum of Phred-scaled base quality scores for the alternate allele) as a tiebreaker. Complex nucleotide variants that did not alter sequence length were decomposed into individual single nucleotide polymorphisms (SNPs). Variants outside the ROI were discarded. These filters are supported by the benchmark described in "Benchmarking cluster depths".

##### 9. Mutation profiling

The filtered variants were applied to the reference sequence for each cluster. Clusters containing indels were counted separately and excluded from amino-acid analysis. For the remaining clusters, the resulting ROI sequence was translated using the standard genetic code and compared to the reference protein sequence to identify amino-acid substitutions. Synonymous substitutions were ignored, and amino-acid positions were reported in full protein coordinates. Clusters were then grouped by their set of non-synonymous amino-acid substitutions, producing a count table of mutation profiles (e.g., "F751Y M753T", "wild", or "indels") and their corresponding cluster counts. These tables were used as input for downstream analyses.

#### Benchmarking cluster depths

We analyzed eight independent ONT R10.4.1 datasets to empirically determine an appropriate minimum cluster depth for UMI-based consensus generation and estimate error rates post denoising. Our benchmarking approach was adapted from Zahm et al.[7]. A series of subsampled clusters were evaluated against a high-depth reference consensus treated as ground truth. In our case, for each dataset, the 100 highest-depth clusters were selected, with read depths ranging from 283 to 162,321 reads. From each reference cluster, 1,000 subsampled clusters were generated at each tested depth (2-20 and 25 reads) by randomly drawing reads from the original cluster alignments. Variants were called in the ROI of each cluster using FreeBayes (ploidy = 1, --genotype-qualities) using parameters described before. Single-read clusters were excluded from this benchmark as FreeBayes does not output variants for them by default, and they are expected to yield largely spurious variant calls under the AO > RO criterion.

Consensus sequences were reconstructed from the variant calls using "bcftools consensus". For the high-depth reference clusters, the GT = "alt" filter was used and is reliable in haploid mode at high depth. For the subsampled clusters we chose the AO > RO filter used in our analysis. VCF filtering parameters could in principle be optimized per dataset.

Results from all eight datasets were pooled to provide a robust, cross-condition estimate of consensus accuracy. Differences between each subsampled consensus and its corresponding reference consensus were quantified using Levenshtein distances (python-Levenshtein v0.27.3, https://github.com/miohtama/python-Levenshtein), expressed as base-level error rates and converted to Phred-style base-level Q scores (lower bound of the 95% confidence interval computed using a beta distribution).

$$Q = -10 \log_{10}(error\ rate)$$

In parallel, the fraction of fully correct consensuses was computed for each depth.

As shown in **Supplementary Figure 9c,d**, consensus quality improves sharply with cluster depth: depth 2 yields a base-level Q≥26.8 and 67.7% fully correct consensus sequences; depth 3 reaches Q≥35.9 and 96.4% fully correct; by depth 5, Q≥48.7 and 99.6% of consensus sequences are fully correct. Beyond depth 5, the fraction of fully correct consensus sequences shows only marginal gains, from 99.6% to 99.9% at depth 25, while base-level Q scores continue to improve more gradually, reaching Q≥57.6 at depth 25. Consequently, we used a cluster depth of 5 throughout this study.

#### Supplementary figures

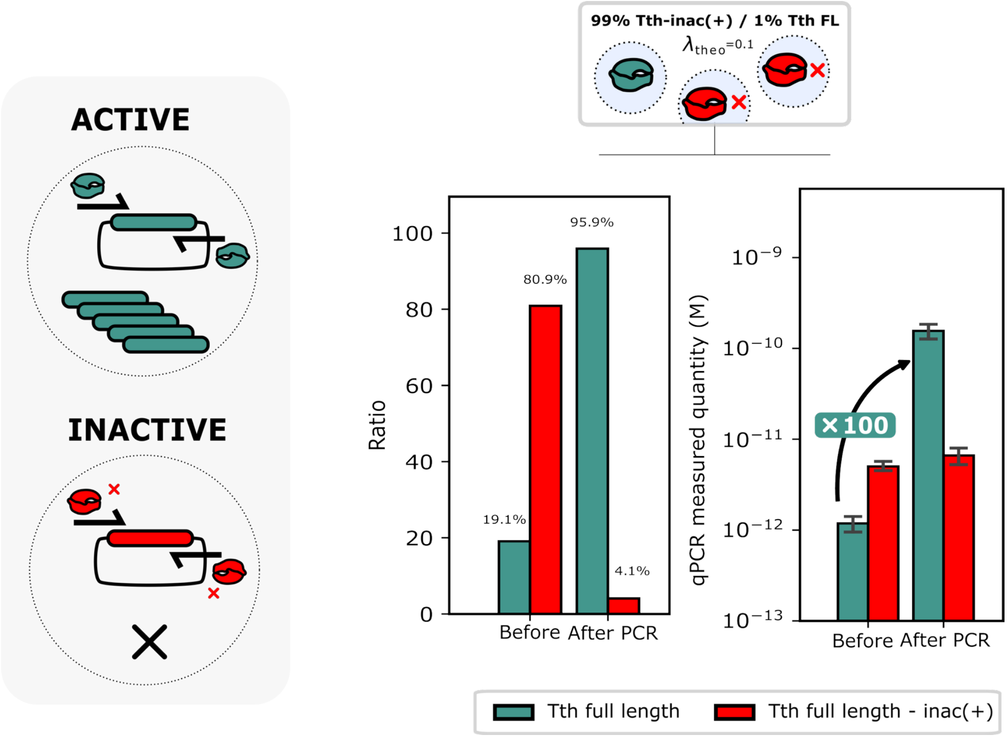

**Supplementary Figure 1 | Proof-of-concept (mock selection) of CSR efficiency implemented in a microfluidic setup.**

A mock selection experiment was performed to assess the efficiency of the Compartmentalized Self-Replication (CSR) workflow using microfluidic library encapsulation. A theoretical Poisson parameter λ of 0.1 was used to achieve mostly single bacterium per drop encapsulation, with a starting population composed in volume of 99% inactive and 1% active polymerases. After one CSR cycle, performed either with or without an emulsion PCR step, the proportion of active and inactive variants was quantified by qPCR. A single selection round reduced the inactive population from 81% to 4%, demonstrating the strong enrichment capacity of the system.

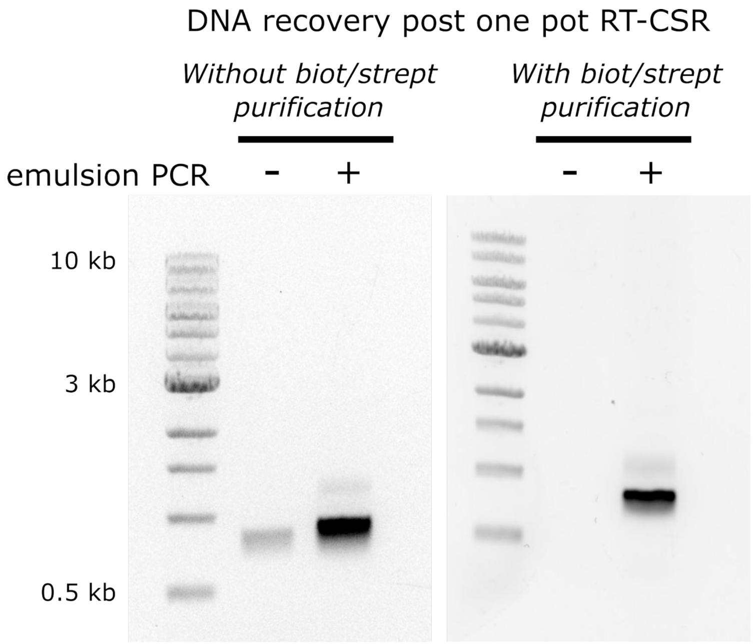

**Supplementary Figure 2 | Biotin-streptavidin purification improves specificity of DNA recovery during CSR.**

A biotin-streptavidin affinity purification step was introduced prior to DNA recovery at the end of the RT-CSR cycle to selectively isolate variants that had successfully reverse-transcribed the modified RNA patch. After RT-CSR, reaction products were incubated with streptavidin-coated magnetic beads (60 min) to capture biotinylated DNA fragments. Beads were washed with NaOH solution (30 mM, 37 °C) to remove unbound or non-specific amplicons. Bound material is then amplified by PCR.

PCR analysis of DNA recovered with or without the biotin-streptavidin purification step revealed that the faint band corresponding to parasitic amplicons, visible in samples not subjected to emulsion PCR, was eliminated after purification. This demonstrates improved specificity and reduced background amplification.

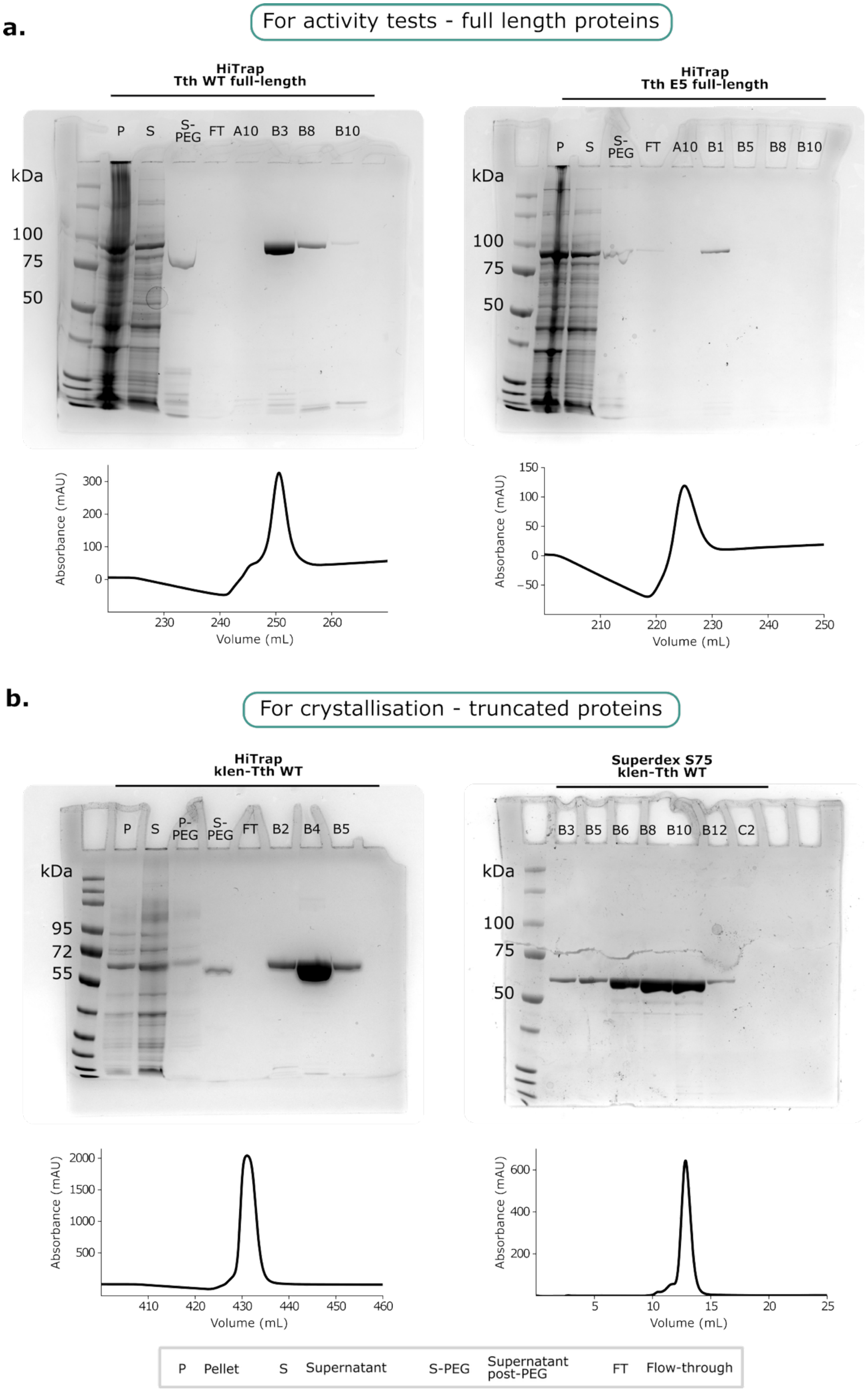
a.
For activity tests - full length proteins
HiTrap
Tth WT full-length
P S S-PEG FT A10 B3 B8 B10
kDa
100
75
50
HiTrap
Tth E5 full-length
P S S-PEG FT A10 B1 B5 B8 B10
kDa
100
75
50
Absorbance (mAU)
Volume (mL)
b.
For crystallisation - truncated proteins
HiTrap
klen-Tth WT
P S P-PEG S-PEG FT B2 B4 B5
kDa
95
72
55
Superdex S75
klen-Tth WT
B3 B5 B6 B8 B10 B12 C2
kDa
100
75
50
Absorbance (mAU)
Volume (mL)
P Pellet
S Supernatant
S-PEG Supernatant post-PEG
FT Flow-through

**Supplementary Figure 3 | Chromatograms and Coomassie stained SDS-PAGE gels of purified polymerases (WT Tth and evolved variant E5).**

**(a)** For activity assays, polymerases were expressed as full-length proteins. After PEG precipitation, the supernatant was loaded onto a HiTrap ion-exchange column and eluted with a NaCl gradient. The indicated fractions were analysed by SDS-PAGE: pellet (P), clarified lysate supernatant (S), supernatant after PEG precipitation (S-PEG), and flow-through from the HiTrap column (FT).

**(b)** For crystallization of the Tth WT polymerase, the protein was N-terminally truncated to obtain the equivalent of the Klenow fragment (without the 5'-3' exonuclease N-terminal domain). Following heparin affinity chromatography, the sample was further purified by size-exclusion chromatography on a Superdex 75 column. Relevant fractions from the chromatographic steps were analysed by SDS-PAGE as in panel (a).

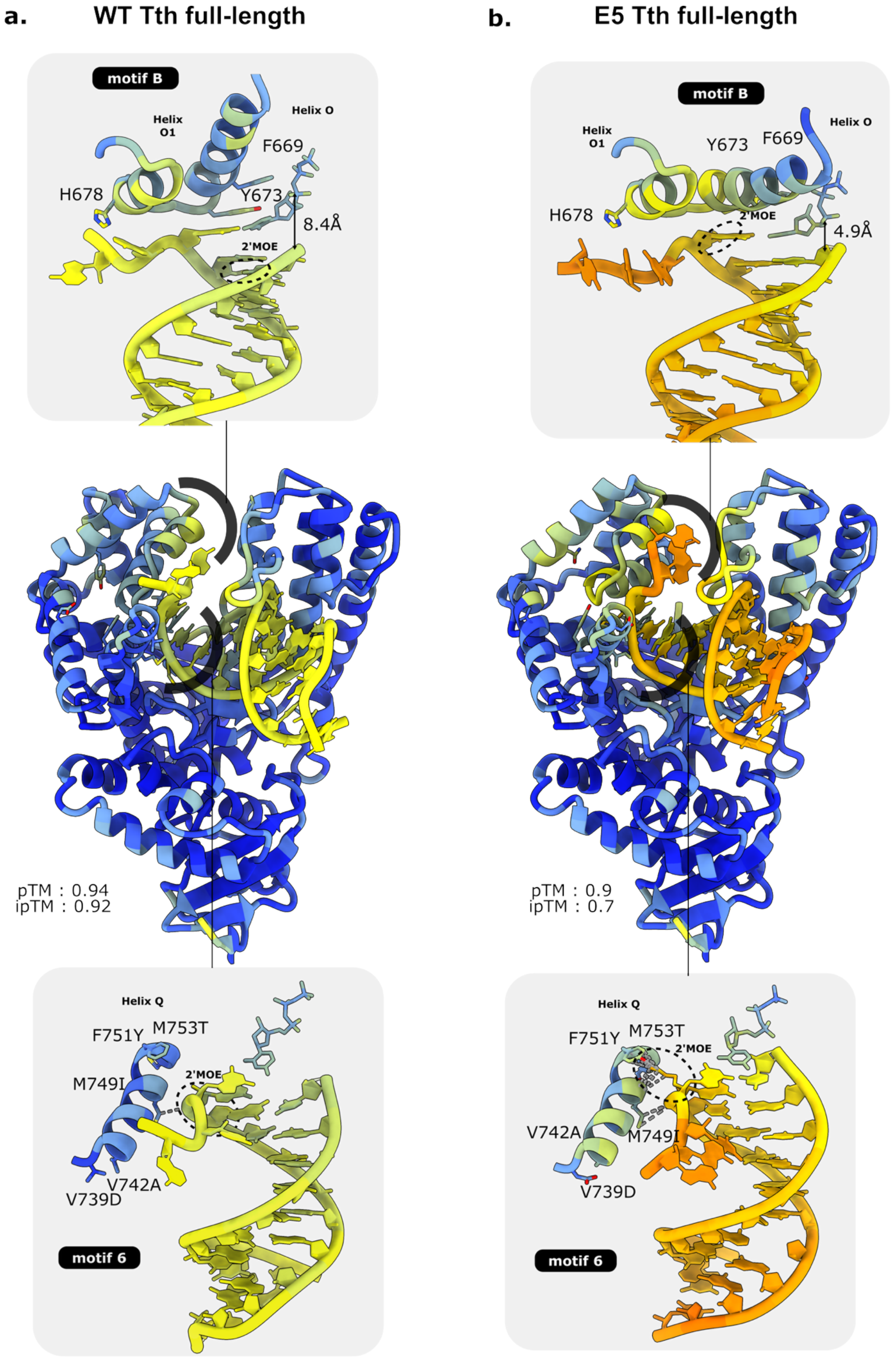
a.
WT Tth full-length
b.
E5 Tth full-length
motif B
Helix O1
Helix O
F669
H678
Y673
8.4Å
2'MOE
motif B
Helix O1
Y673
F669
Helix O
H678
2'MOE
4.9Å
pTM : 0.94
ipTM : 0.92
pTM : 0.9
ipTM : 0.7
Helix Q
F751Y
M753T
M749I
2'MOE
V742A
V739D
motif 6
Helix Q
F751Y
M753T
2'MOE
V742A
M749I
V739D
motif 6

**Supplementary Figure 4 | Structural basis predictions for enhanced 2'MOE template processing by evolved polymerase E5.**

AlphaFold3 predictions of polymerase complexes with RNA substrate bearing 2'MOE modification at the templating position plus incoming nucleotide. The most frequently observed conformation (4 of 5 models) is shown. Both predictions achieved high confidence scores (pTM > 0.9, ipTM > 0.7) with coherent duplex geometry and proper base pairing. Structures are coloured by pLDDT. **(a)** WT Tth DNA polymerase structure and **(b)** evolved variant E5 structures are shown at three magnifications. **Top panel**, motif B (with helices O and O1) in closed conformation enabling proper positioning of the 2'MOE-modified n+1 templating nucleotide and incoming nucleotide for E5. In WT, motif B adopts an open conformation causing the incoming nucleotide to mis-pair with the n+2 base, resulting too far 3'-OH positioning (8.4 Å from α-phosphate). In contrast, E5's closed motif B conformation maintains proper 3'-OH distance (4.9 Å), enabling catalysis. **Middle panel**, overview of full enzyme architecture with modified RNA duplex, finger, palm, and thumb domains. **Bottom panel**, helix Q (motif 6) showing or not E5 mutations. F751Y mutation interacts strongly to stabilize the template 2'MOE modification.

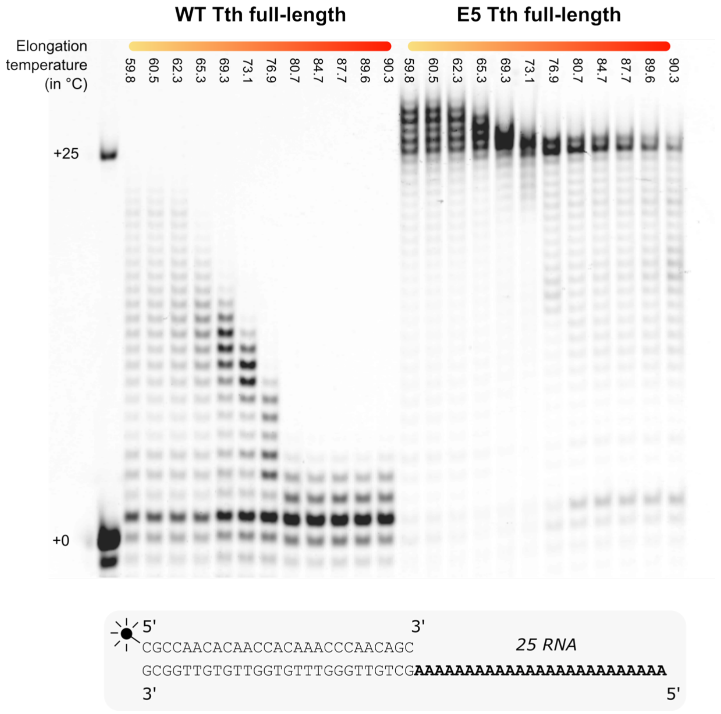

**Supplementary Figure 5 | Temperature dependence of reverse transcriptase activity.**

Reverse transcriptase activity was tested on a 25-nucleotide RNA substrate across a 60 to 90 °C temperature range. The E5 variant fully converts the substrate to complementary DNA across the entire range, whereas the WT enzyme becomes inactive above 80 °C and never achieves complete RNA to cDNA conversion.

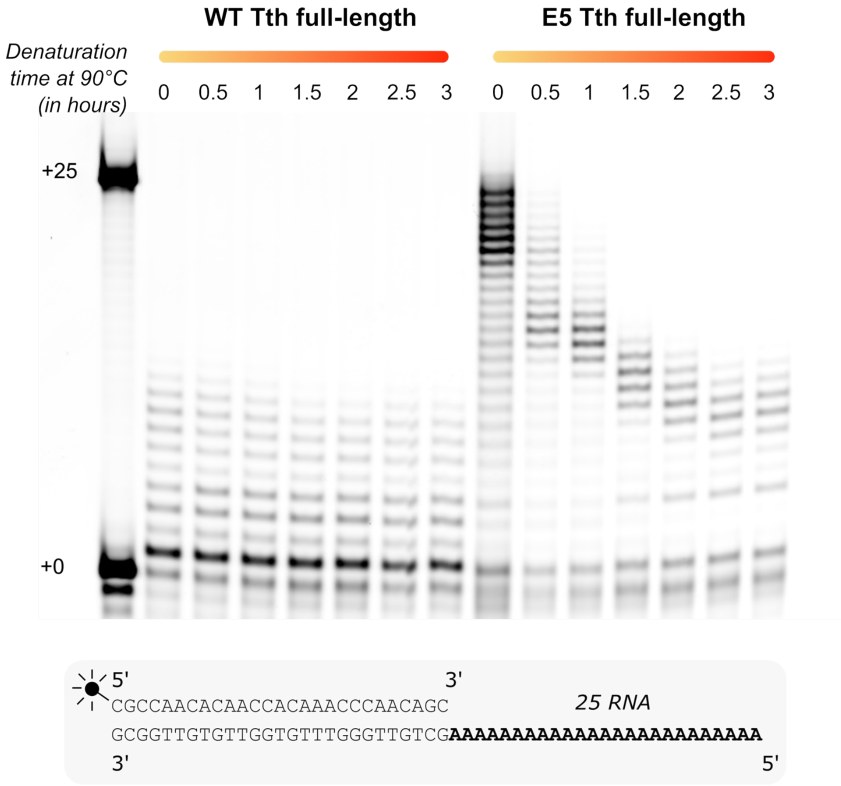

**Supplementary Figure 6 | E5 reverse transcriptase activity after heat stress.**

WT Tth polymerase and evolved variant E5 were subjected to thermal denaturation at 90 °C for up to 3 hours, followed by reverse transcriptase activity assays (30 min, 72 °C). The WT enzyme showed low basal activity at any time point, whereas E5 retained reverse transcriptase activity even after 90 minutes of heat treatment at 90 °C.

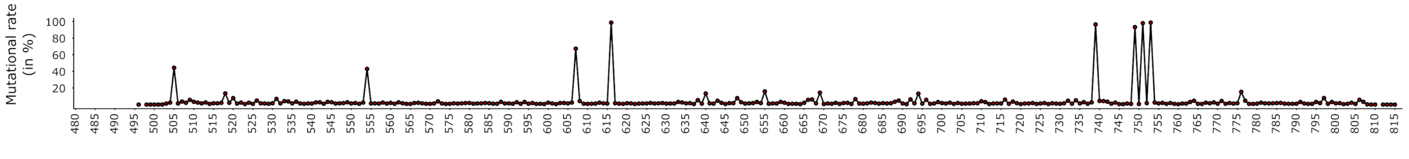

**Supplementary Figure 7 | Mutational hotspots in the final IE013 library.**

Mutation frequencies along the mutated region at the end of directed evolution. Positions 505, 554, 607, 616, 739, 749, 751, and 753 stand out, exhibiting mutation rates exceeding 20% with 616, 739, 749,751 and 753 close to 100%.

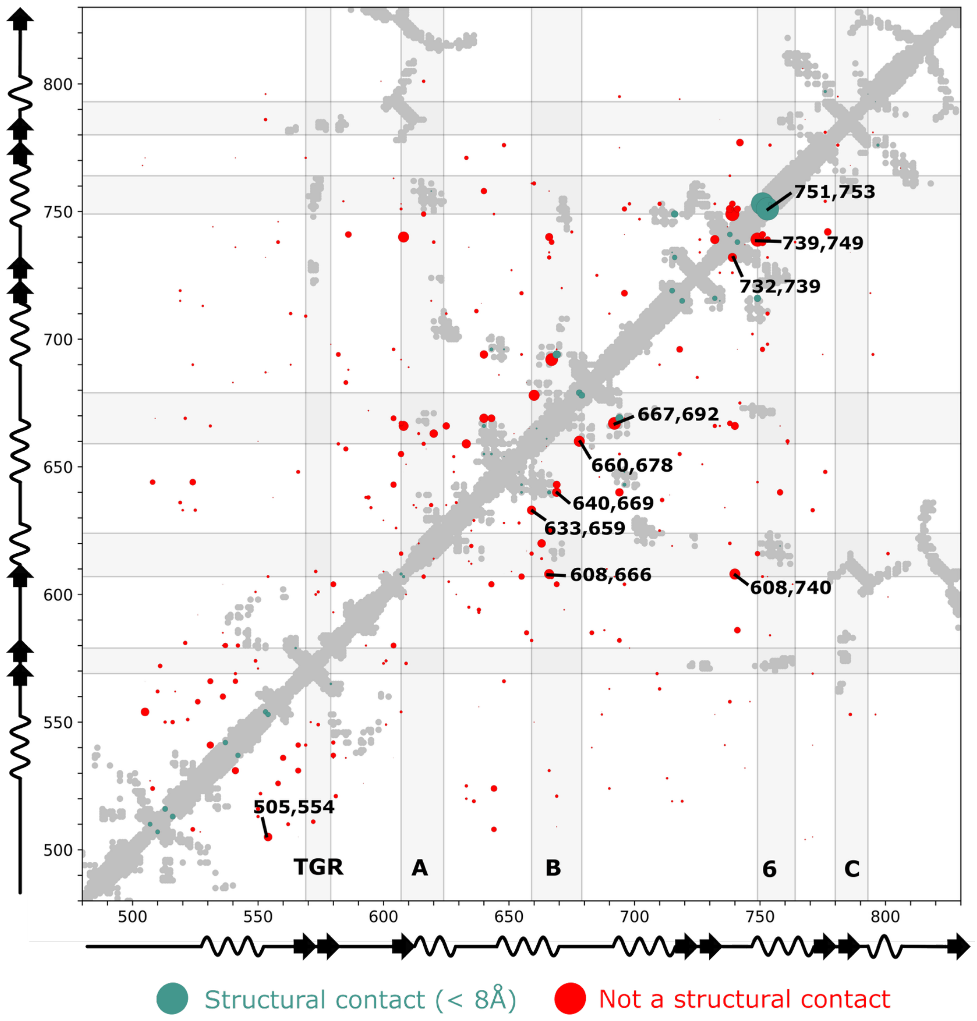

**Supplementary Figure 8 | Evolutionary couplings identified by mfDCA analysis and their structural relevance.**

Couplings identified by mean-field Direct Coupling Analysis (mfDCA) were aggregated across directed evolution rounds to highlight those that have been most conserved or influential during evolution. These aggregated couplings were then compared with structural data to determine whether functionally important contacts also correspond to physical contacts (Cα-Cα distance < 8 Å). The 10 stronger couplings are annotated. Most of the identified couplings are not structural, and the most significant ones predominantly involve motif B and motif 6, suggesting their key role in acquisition of XNA-RT activity.

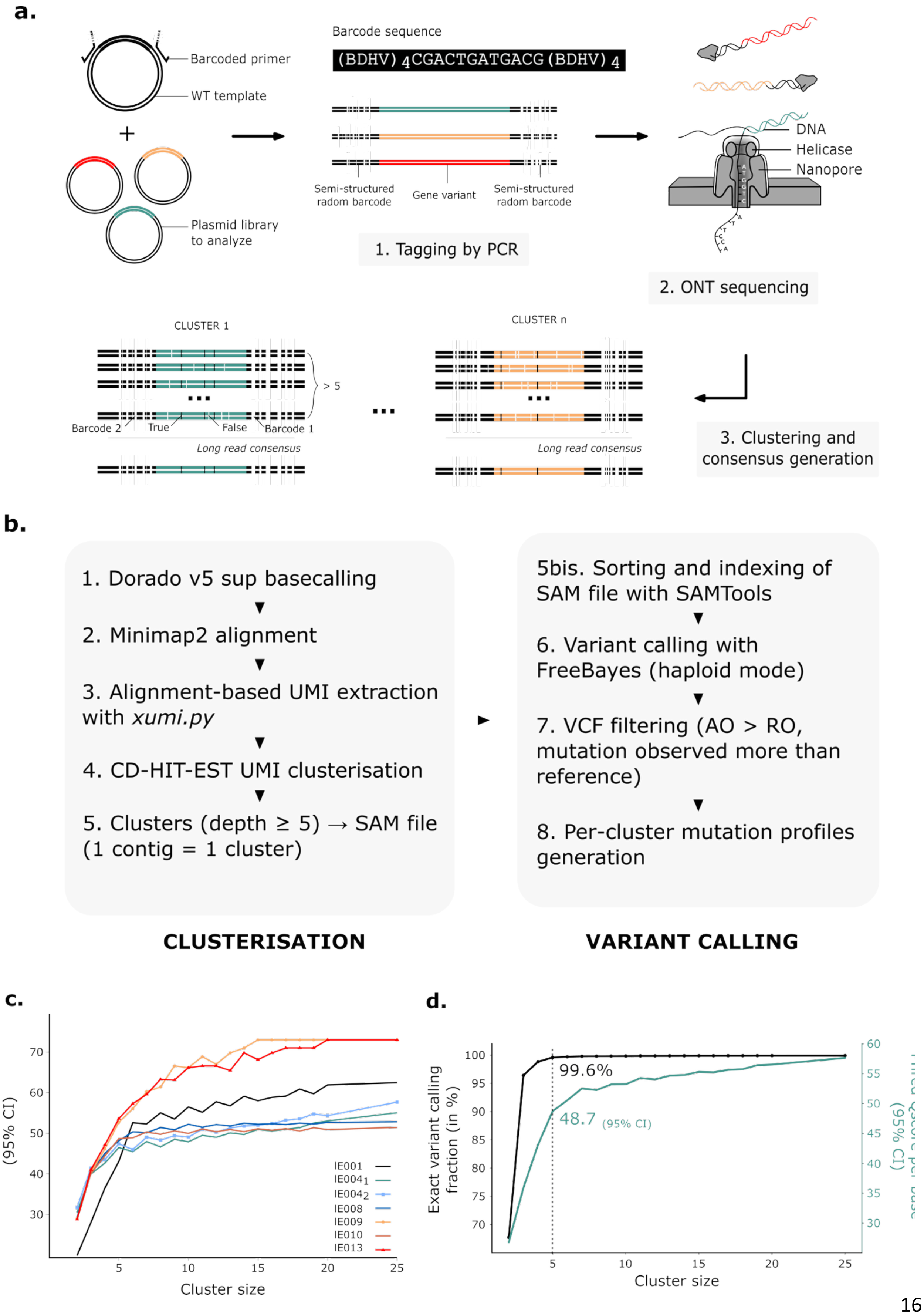

a.
Barcoded primer
WT template
+
Plasmid library
to analyze
Barcode sequence
(BDHV) 4CGACTGATGACG (BDHV) 4
Semi-structured
radom barcode
Gene variant
Semi-structured
radom barcode
1. Tagging by PCR
DNA
Helicase
Nanopore
2. ONT sequencing
CLUSTER 1
> 5
Barcode 2
True
False
Barcode 1
Long read consensus
CLUSTER n
Long read consensus
3. Clustering and
consensus generation
b.
1. Dorado v5 sup basecalling
2. Minimap2 alignment
3. Alignment-based UMI extraction
with xumi.py
4. CD-HIT-EST UMI clusterisation
5. Clusters (depth ≥ 5) → SAM file
(1 contig = 1 cluster)
5bis. Sorting and indexing of
SAM file with SAMTools
6. Variant calling with
FreeBayes (haploid mode)
7. VCF filtering (AO > RO,
mutation observed more than
reference)
8. Per-cluster mutation profiles
generation
CLUSTERISATION
VARIANT CALLING
c.
(95% CI)
Cluster size
IE001
IE004₁
IE004₂
IE008
IE009
IE010
IE013
d.
Exact variant calling
fraction (in %)
99.6%
48.7 (95% CI)
(95% CI)
Cluster size

**Supplementary Figure 9 | Sequencing and bioinformatic denoising pipeline for accurate variant identification.**

**(a)** The experimental sequencing pipeline is based on tagging of the target region using semi-structured 44-base unique molecular identifiers (UMI). After cloning, the UMI-flanked region of interest was amplified by PCR and sequenced using Oxford Nanopore Technologies (ONT) on a MinION flow cell (Ligation Sequencing of Amplicons, kit V14). Following 24 hours of sequencing, raw signals were basecalled with the latest Dorado model (SUP v5.0.0). The resulting reads were processed through a bioinformatic denoising workflow combining clustering and consensus sequence derivation to separate true mutations from sequencing noise.

**(b)** The denoising and variant-calling pipeline begins with high-accuracy basecalling (model: dna_r10.4.1_e8.2_400bps_sup@v5.0.0). Reads are aligned to the Tth WT reference sequence using Minimap2. Barcodes are extracted and reads are clustered with CD-HIT-EST to group reads originating from the same molecule. Only clusters containing at least five reads are retained. Within each cluster, variants are called using FreeBayes and filtered based on variant quality metrics to exclude sequencing artefacts, retaining only true mutations.

**(c)** Per-dataset empirical Phred Q scores are plotted as a function of cluster depth. Base-level Q scores (95% CI lower bounds) are computed from 1,000 subsampled clusters per depth across 100 high-depth reference clusters, shown separately for each of seven independent datasets. Consensus sequences were reconstructed using BCFtools consensus with the AO > RO filter applied to FreeBayes variant calls.

**(d)** Fraction of fully correct consensus sequences **(left axis,** black**)** and base-level Phred Q scores (95% CI lower bounds, **right axis**, green) are computed from 1,000 subsampled clusters per depth across 100 high-depth reference clusters all pooled from eight independent datasets. Consensus sequences were reconstructed as in **(c)**. Dashed vertical line indicates the ≥5 depth threshold used in the main pipeline.

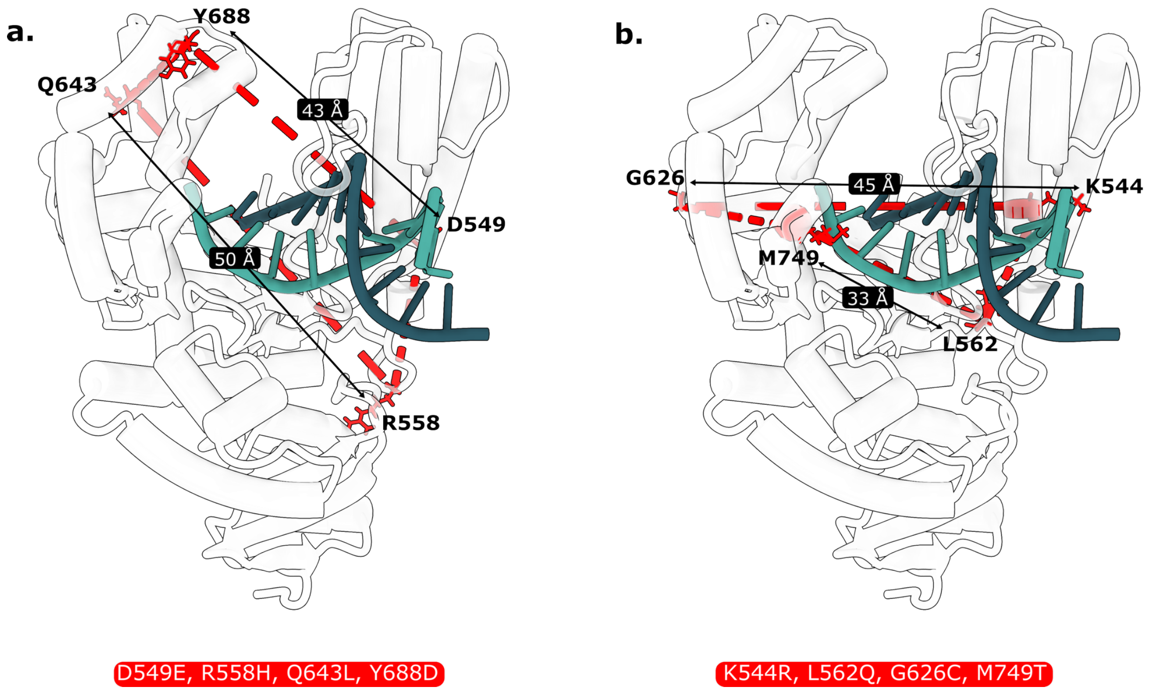

**Supplementary Figure 10 | Four-body couplings in the Tth DNA polymerase I.**

Analysis of the fitness landscape using the protein language model ESM2-3B reveals the presence of higher-order (four-body) couplings within the Tth polymerase A. The identified couplings connect residues located in the finger and thumb domains, which are spatially distant, highlighting strong long-range cooperativity and conformational dynamics within the enzyme.

Two representative examples are shown: **(a)** the set of coupled mutations D549E, R558H, Q643L, and Y688D includes a maximal Cα-Cα distance of 50 Å between Q643 and Y688; **(b)** similarly, the set K544R, L562Q, G626C, and M749T exhibits a maximal distance of 45 Å between G626 and K544. These findings demonstrate the ability of protein language models to detect atomically precise, long-range epistatic interactions that are difficult to capture by conventional approaches.

### Supplementary tables

| | KlenTth_2'MOE/RNA_template |
|---|---|
| **Data collection** | |
| Space group | $P2_12_12_1$ |
| a. b. c (Å) | 80.17 94.94 238.62 |
| α, β, γ (◦) | 90.0 90.0 90.0 |
| Wavelength (Å) | 0.97856 |
| Resolution (Å) | 119.3– 1.924 (2.209-1.924)* |
| Staraniso resolution limit (Å) | 2.752 2.371 1.930 |
| R-pim | 0.078 (0.629) |
| Completeness (%) | 94.1 (72.0) |
| Multiplicity | 8.5 (7.7) |
| I/σ(I) | 7.4 (1.6) |
| CC1/2 | 0.966 (0.638) |
| **Refinement** | |
| No. of reflections | 75 007 (4 787) |
| Rwork / Rfree | 0.194/0.229 (0.283/0.309) |
| No. non H atoms | 10 617 |
| No amino acids | 1 076 |
| No Ligands | 44 |
| No Solvent | 1311 |
| **Protein geometry** | |
| Molprobity score | 1.19 |
| RMSD—bonds (Å) | 0.006 |
| RMSD—angles (◦) | 0.755 |
| Ramachandran favored (%) | 97.48 % |
| Ramachandran outliers (%) | 0.19 % |
| Poor rotamers (%) | 0.23 % |
| Clashscore | 2.91 |
| **B-factors ($Å^2$)** | |
| Mean B-factor | 32.1 |
| Macromolecules | 31.0 |
| Ligands | 39.2 |
| Solvent | 35.7 |

*Figures in parenthesis represent values on the highest resolution shell

**Supplementary Table 1|Data collection and refinement statistics for KlenTth WT with 2'MOE/RNA template.**

| Name | Use | Sequence |
|---|---|---|
| **iE029** | epPCR | ATTATGGATCTAGAGACTAACGGAGCTATACGXXGCTGGCAAGTGTAGCGGTCAC |
| **iE030** | epPCR | ATATAGTATCGTTCTCGGTATATGGTGCATCGXXATTAGGAAGCAGCCCAGTAGTAGG |
| **iE046** | Insert cloning domain 500 bp | TCAGCGTATTCGTCGTGCCTTTG |
| **iE047** | Insert cloning domain 500 bp | GGCCAGTTTCATCAGATCGGCTGC |
| **iE048** | Vector cloning domain 500 bp | CAGCCGATCTGATGAAACTGGCC |
| **iE049** | Vector cloning domain 500 bp | AAGGCACGACGAATACGCTGACC |
| **iE092** | Insert cloning domain 1000 bp | GGCCAGAGGATATGCTTTTTCCATTG |
| **iE101** | Insert cloning domain 1000 bp | GTGCTGTTTGATGAACTGCGTCTG |
| **iE102** | Vector cloning domain 1000 bp | ACGCAGTTCATCAAACAGCACACG |
| **iE090** | Vector cloning domain 1000 bp | TGGAAAAAGCATATCCTCTGGCCGTTC |
| **iE033** | Selective pressure 1 2’MOE/1RNA | ATTATGGATCTAGAGACTAACGGAGCTATACG<A>(A)GCTGGCAAGTGTAGCGGTCAC |
| **iE034** | Selective pressure 1 2’MOE/1RNA | ATATAGTATCGTTCTCGGTATATGGTGCATCG<T>(U)ATTAGGAAGCAGCCCAGTAGTAGG |
| **iE035** | Selective pressure 2 2’MOE/2RNA | ATTATGGATCTAGAGACTAACGGAGCTATACG<G>(A)<A>(A)GCTGGCAAGTGTAGCGGTCAC |
| **iE036** | Selective pressure 2 2’MOE/2RNA | ATATAGTATCGTTCTCGGTATATGGTGCATCG<T>(A)<C>(A)ATTAGGAAGCAGCCCAGTAGTAGG |
| **iE104** | Outnested primer biotinylated | [BIOTEG]AAAAAAAAAAATCGTTCTCGGTATATGGTGCATCG |
| **iE105** | Outnested primer biotinylated | [BIOTEG]AAAAAAAAAATGGATCTAGAGACTAACGGAGCTATACG |
| **iE124** | Primer kinetics | [ATTO488]c*g*c*caacacaaccacaaaccccaac*a*g*c |
| **iE135** | Ladder kinetics +4 | [ATTO488]c*g*c*caacacaaccacaaaccccaac*a*g*cTTTT |
| **iE144** | Ladder kinetics +10 | [ATTO488]c*g*c*caacacaaccacaaaccccaac*a*g*cTTTTGCGCGC |
| **iE136** | Template kinetics DNA | GCGCGCAAAAGCTGTTGGGGTTTGTGGTTGTGTTG*G*C*G |
| **iE137** | Template kinetics 1 RNA | GCGCGC(A)AAAGCTGTTGGGGTTTGTGGTTGTGTTG*G*C*G |
| **iE138** | Template kinetics 1 2’OMe | GCGCGC[2OMeA]AAAGCTGTTGGGGTTTGTGGTTGTGTTG*G*C*G |
| **iE139** | Template kinetics 1 2’MOE | GCGCGC<A>AAAGCTGTTGGGGTTTGTGGTTGTGTTG*G*C*G |
| **iE111** | Primer RT consecutive | [ATTO647N]C*G*C*CAACACAACCACAAACCCCAAC*A*G*C |
| **iE109** | Template DNA consecutive | AAAAAAAAAAAAAAAAAAAAAAAAAGCTGTTGGGGTTTGTGGTTGTGTTG*G*C*G |
| **iE114** | Template RNA consecutive | (A)(A)(A)(A)(A)(A)(A)(A)(A)(A)(A)(A)(A)(A)(A)(A)(A)(A)(A)(A)(A)(A)(A)(A)(A)GCTGTTGGGGTTTGTGGTTGTGTTG*G*C*G |
| **iE134** | Template 2’OMe consecutive | [2OMeA][2OMeA][2OMeA][2OMeA][2OMeA][2OMeA][2OMeA][2OMeA][2OMeA][2OMeA][2OMeA][2OMeA][2OMeA][2OMeA][2OMeA][2OMeA][2OMeA][2OMeA][2OMeA][2OMeA][2OMeA][2OMeA][2OMeA][2OMeA][2OMeA]GCTGTTGGGGTTTGTGGTTGTGTTG*G*C*G |
| **iE161** | Template RT 2’OMe - RNA (1/4) | (A)[2OMeA](A)(A)(A)[2OMeA](A)(A)(A)[2OMeA](A)(A)(A)[2OMeA](A)(A)(A)[2OMeA](A)(A)(A)[2OMeA](A)(A)(A)GCTGTTGGGGTTTGTGGTTGTGTTG*G*C*G |

| | | |
|---|---|---|
| **iE162** | Template RT 2’OMe - RNA (1/2) | (A)[2OMeA](A)[2OMeA](A)[2OMeA](A)[2OMeA](A)[2OMeA](A)[2OMeA](A)[2OMeA](A)[2OMeA](A)[2OMeA](A)[2OMeA](A)[2OMeA](A)[2OMeA](A)GCTGTTGGGGTTTGTGGTTGTGTTG*G*C*G |
| **iE133** | Template mixed consecutive | (A)<A>(A)<A>(A)<A>(A)<A>(A)<A>(A)<A>(A)<A>(A)<A>(A)<A>(A)<A>(A)<A>(A)<A>(A)GCTGTTGGGGTTTGTGGTTGTGTTG*G*C*G |
| **iE108** | KlenTth truncation | CTGGAAGAAGCCCCTTGGC |
| **iE008** | KlenTth truncation | CATTGAAGTGTATATCTCCTTCTTAAAGTTAAACAAAATTATTTCTAGAG |
| **iE087** | Library barcoding | TGCTGCTCAGGCGACCBDHVBDHVBDHVBDHVCGACTGATGACGBDHVBDHVBDHVBDHVTACGCGGATGAACCAGGCTCG |
| **iE088** | Library barcoding | ATGGAAAAAGCATATCCTCTGGCCBDHVBDHVBDHVBDHVCGACTGATGACGBDHVBDHVBDHVBDHVCGCAAAAGGTTAAGGTGGCTCAG |
| **iE121** | Library barcoding 1000bp | GACGCAGTTCATCAAACAGCACACBDHVBDHVBDHVBDHVCGACTGATGACGBDHVBDHVBDHVBDHVAGACGAAAAACTTCTTCTTCTAAACGACG |
| **iE106** | Xtallo primer | GACCACGGCGC |
| **iE107** | Xtallo template | AAA<G>(G)GCGCCGTGGTC |

**Supplementary Table 2| Primers and templates for all polymerase studies, mutagenesis and directed evolution.**

Different nucleotide chemistries are denoted as follows: plain text for DNA; (X) for RNA; [2OMeX] for 2’OMe RNA; and ⟨X⟩ for 2’MOE RNA.

| Library | Number of raw reads | Number of clusterized reads (clust. size >5) | Number of clusters (= denoised reads) | Number of unique variants | Number of unique variants with no stop | Share of stop into population (in %) | Mean size of clusters |
|---|---|---|---|---|---|---|---|
| **lE001** | 5.9 M | 4.6 M | 43 296 | 16 587 | 15 075 | 9% | 106 |
| **lE004$_1$** | 4.5 M | 3.8 M | 24 532 | 1 623 | 1 549 | 5% | 155 |
| **lE004$_2$** | 3.8 M | 2.5 M | 15 102 | 1 558 | 1 489 | 4% | 164 |
| **lE008** | 6.3 M | 4.2 M | 97 255 | 60 943 | 49 687 | 18% | 43 |
| **lE009** | 5.1 M | 3.5 M | 83 602 | 52 551 | 45 647 | 13% | 42 |
| **lE010** | 12 M | 4.4 M | 66 106 | 38 254 | 32 880 | 14% | 66 |
| **lE013** | 6.3 M | 5.0 M | 91 547 | 58 284 | 50 457 | 13% | 55 |
| **Total** | **44 M** | **28 M** | **421 k** | **230 k** | **196 k** | | |

**Supplementary Table 3| Summary of the high-throughput sequencing (MinION Oxford Nanopore Technology) experiments performed throughout the evolution campaign.**

For each library, the table lists the library name, the number of raw reads before clustering, the number of reads aligned to the reference, the number of clusters (defined as unique barcode pairs supported by ≥5 reads) and their mean cluster size, the number of unique variants recovered, the number of unique variants without premature stop codons and the fraction of reads carrying stop codons in the library.
